# Cell-specific crosstalk proteomics reveals cathepsin B signaling as a driver of glioblastoma malignancy near the subventricular zone

**DOI:** 10.1101/2023.08.19.553966

**Authors:** Emily S. Norton, Lauren A. Whaley, Vanessa K. Jones, Mieu M. Brooks, Marissa N. Russo, Dmytro Morderer, Erik Jessen, Paula Schiapparelli, Andres Ramos-Fresnedo, Natanael Zarco, Anna Carrano, Wilfried Rossoll, Yan W. Asmann, TuKiet T. Lam, Kaisorn L. Chaichana, Panos Z. Anastasiadis, Alfredo Quiñones-Hinojosa, Hugo Guerrero-Cázares

**Affiliations:** Department of Neurosurgery, Mayo Clinic, Jacksonville, FL 32224, USA; Neuroscience Graduate Program, Mayo Clinic Graduate School of Biomedical Sciences, Mayo Clinic, Jacksonville, FL 32224, USA; Regenerative Sciences Training Program, Mayo Clinic Graduate School of Biomedical Sciences, Mayo Clinic, Jacksonville, FL 32224, USA; Department of Biology, University of North Florida, Jacksonville, FL 32224, USA; Department of Neuroscience, Mayo Clinic, Jacksonville, FL 32224, USA; Division of Biomedical Statistics and Informatics, Department of Health Sciences Research, Mayo Clinic, Jacksonville, FL 32224, USA; Keck MS and Proteomics Resource, Yale School of Medicine, New Haven, CT 06510, USA; Department of Molecular Biophysics and Biochemistry, Yale School of Medicine, New Haven, CT 06510, USA; Department of Cancer Biology, Mayo Clinic, Jacksonville, FL 32224, USA

**Author notes:** Corresponding author and lead contact, Hugo Guerrero-Cázares., Twitter: @HugoGuerreroLab.

**Keywords:** brain cancer, lateral ventricle, stem cell niche, neurogenesis, glioma, subventricular zone, MetRS L274G, proteomics, click chemistry, senescence

## Abstract

Glioblastoma (GBM) is the most prevalent and aggressive malignant primary brain tumor. GBM proximal to the lateral ventricles (LVs) is more aggressive, potentially due to subventricular zone (SVZ) contact. Despite this, crosstalk between GBM and neural stem/progenitor cells (NSC/NPCs) is not well understood. Using cell-specific proteomics, we show that LV-proximal GBM prevents neuronal maturation of NSCs through induction of senescence. Additionally, GBM brain tumor initiating cells (BTICs) increase expression of CTSB upon interaction with NPCs. Lentiviral knockdown and recombinant protein experiments reveal both cell-intrinsic and soluble CTSB promote malignancy-associated phenotypes in BTICs. Soluble CTSB stalls neuronal maturation in NPCs while promoting senescence, providing a link between LV-tumor proximity and neurogenesis disruption. Finally, we show LV-proximal CTSB upregulation in patients, showing the relevance of this crosstalk in human GBM biology. These results demonstrate the value of proteomic analysis in tumor microenvironment research and provide direction for new therapeutic strategies in GBM.

**Highlights:**

- Periventricular GBM is more malignant and disrupts neurogenesis in a rodent model.
- Cell-specific proteomics elucidates tumor-promoting crosstalk between GBM and NPCs.
- NPCs induce upregulated CTSB expression in GBM, promoting tumor progression.
- GBM stalls neurogenesis and promotes NPC senescence via CTSB.

## Introduction

Glioblastoma (GBM) is the most common and aggressive primary brain tumor in adults^1^. The location of GBM contributes to patient outcomes, where tumors in contact with the lateral ventricles (LVs) result in increased tumor expression of stem cell genes, increased incidence of distal recurrence, and decreased median overall survival independent of age and extent of resection^2–9^. This may be due to the presence of the subventricular zone (SVZ), the largest neurogenic niche in mammals, which is located along the lateral wall of the LVs^10–12^.

The SVZ contains populations of neural stem and progenitor cells (NSC and NPCs, respectively) throughout life, though the human SVZ lacks the migrating neuroblast chains observed in rodent models^11,12^. NSCs extracted from the adult SVZ have been characterized in vitro by their ability to proliferate, self-renew, and differentiate into multiple lineages^13–17^. Interestingly, there is a subpopulation of stem-like GBM cells located throughout the tumor, termed brain tumor initiating cells (BTICs), with very similar properties to NSCs. BTICs are able to form neurospheres, differentiate into multiple progeny, and share several migratory behaviors and markers with NSCs, with the additional ability to form tumors in vivo^18–20^. The presence of these stem-like cells contributes to tumor progression, therapeutic resistance, and worse patient outcome^21^. The biological similarity of these cell types to NSCs supports the idea that stem cell-promoting factors in the SVZ may promote the stemness and progression of nearby GBM tumors, and that NSCs may play a role in GBM malignancy. Indeed, previous work has shown that cell types within the SVZ, both neurogenic and non-neurogenic, are altered in response to LV-associated GBM in rodents^22–24^.

Gene expression in both GBM and the SVZ have been well characterized at a transcriptomic level using combinations of bulk and single cell RNA sequencing^25–32^. Despite this, proteomic characterization is still in earlier stages for these niches^33–37^, and cell-specific proteomics has not yet been performed to differentiate neurogenic and non-neurogenic proteomes. The proteomic evaluation of NPCs as well as GBM BTICs is incredibly important, as studies have shown there is considerable disconnect between the transcriptome and translatome in both the neurogenic cell population of the adult SVZ and the neoplastic cells of GBM tumors. In the SVZ, this is due in large part to differentiation being post-transcriptionally controlled in NPC and their progeny^38^. In GBM, it was concluded that proteomic and phosphoproteomic analysis in addition to transcriptomics is crucial to fully understand malignant pathways at work^37^, again pointing to the necessity of advancing proteomic knowledge of this disease.

In a recent study using a rodent model, we have demonstrated that LV-proximal GBM tumors exhibit faster progression compared to their LV-distal counterparts, mirroring the findings observed in patients. Additionally, we found that these tumors disrupt SVZ neurogenesis^24^. However, it is still unknown what GBM and NPC-specific mechanisms are at play in this bidirectional cellular interaction. Here, we employ a nascent proteomic labeling system via the L274G mutation of methionine tRNA synthetase (MetRS L274G, MetRS*) in order to investigate cell-specific proteomic changes in BTICs and SVZ NPCs when GBM is located in proximity to the LV. We demonstrate that SVZ NSC/NPC neuronal maturation is suppressed, and senescence is promoted in the presence of nearby GBM tumors, which is partially driven by increased cathepsin B (CTSB) expression in tumor cells, implicating the contribution of this signaling axis to the worse prognosis of LV-proximal GBM.

## Results

### Glioblastoma proximity to the lateral ventricles increases tumor malignancy and disrupts normal neurogenic processes

Previous studies have determined that GBM in contact with the LVs is more aggressive than LV-distant counterparts in patients and animal models, potentially due to contact with components of the SVZ^2,8,9,23,24,39–42^. We first investigated whether GBM tumor malignancy was increased and resulted in altered neurogenesis when tumors were close to the LVs in an immunocompetent rodent model. To this end, we generated transgenic mice to express GFP and MetRS* under the Nestin promoter with tamoxifen (TAM)-inducible activation of Cre (Nestin-Cre^ERT2^; STOPflox R26-GFP MetRS L274G (Nes-MetRS* mice), allowing for fate tracking of NSCs. Following TAM administration, mCherry and luciferase (mCh-luc)-positive Gl261 murine glioma cells were implanted at locations proximal and distal to the LV for immunohistochemical and survival analysis. A LV-proximal vehicle (PBS) injected group was included to account for the effects of intracranial injection on the SVZ (**Figure 1A**). Within the tumor, there was an increase in the percent of proliferating pHH3+/mCh+ cells in LV-proximal GBM compared to LV-distal GBM (**Figure 1B-C**). There was also an increased percentage of Sox2+/mCh+ cells (**Figure 1D**) and a decrease in the percentage of Olig2+/mCh+ cells (**Figure 1E**) in LV-proximal tumors, suggesting a more stem-like state of the tumor associated with worse patient outcome. Additionally, there was significantly decreased survival in LV-proximal GBM animals compared to LV-distal GBM counterparts (**Figure 1F**), indicating that this model captures the worse prognosis of LV-proximal GBM observed in patients.

**Figure 1.**
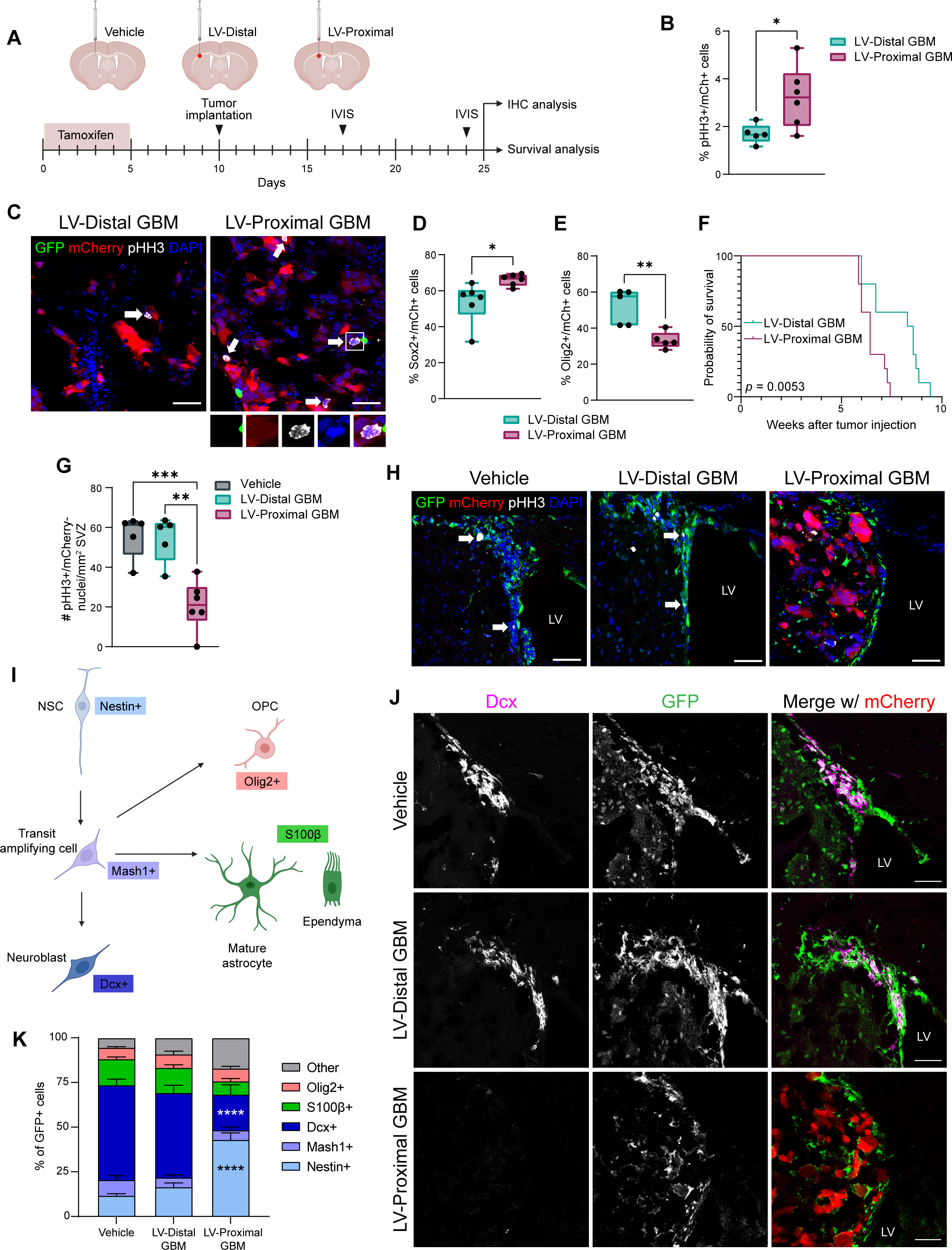
Tumor malignancy is increased and SVZ neurogenesis is decreased in LV-proximal GBM bearing animals. (A) Schematic illustrating the experimental procedure and timeline for Figure 1. (B) Percentage quantification of proliferation in murine GBM cells *in vivo* (n = 5-6) (C) Representative IHC showing increased proliferation in LV-proximal mCh+ tumors. White arrows and inset indicate proliferating tumor cells. Scale bar = 50 µm. (D) Percentage quantification of Sox2+/mCh+ tumor cells (n = 6) (E) Percentage quantification of Olig2+/mCh+ tumor cells (n = 5) (F) Kaplan-Meier curves of survival outcomes for LV-distal GBM and LV-proximal GBM-bearing animals (n = 10) (G) Quantification of proliferative (pHH3+) cells/mm^2^ SVZ (n = 5-6) (H) Representative IHC showing decreased proliferation in the SVZ of the LV-proximal GBM group. White arrows indicate proliferating GFP+ cells. LV = lateral ventricle. Scale bar = 50 µm. (I) Schematic describing neurogenesis markers used and the cell types labeled. (J) Representative IHC showing decreased Dcx+/GFP+ cells in the SVZ of LV-proximal GBM group. LV = lateral ventricle. Scale bar = 50 µm. (K) Percentage quantifications of GFP+ cells in the SVZ that are Nestin+ (light blue), Mash1+ (medium blue), Dcx+ (dark blue), S100b+ (green), and Olig2+ (pink). p < 0.05 *, p < 0.01 **, p < 0.001 ***, p < 0.0001 ****

Conversely, proliferation in the SVZ (pHH3+/mCh-) was decreased in the LV-proximal GBM animals compared to LV-distal GBM animals and to LV-proximal vehicle controls (**Figure 1G-H**). We then examined the phenotype of SVZ GFP+ cells across our conditions using well-studied markers of neurogenesis and glial differentiation to determine cell fate when exiting the Nestin+ stem cell state (**Figure 1I**). We identified a significant decrease in the percentage of GFP+ cells forming neuroblasts (Dcx+/GFP+; **Figure 1J-K**) and a significant increase in the percentage of Nestin+/GFP+ cells (**Figure 1K**) in the SVZ of the LV-proximal GBM group. There were no significant differences in the expression of other cell lineage markers between groups. These results indicate reduced SVZ proliferation and arrested NPC as a result of LV-proximal GBM.

To confirm these effects are due to the interaction between GBM and NSC/NPCs rather than an effect from other factors in the tumor microenvironment, we performed treatment of primary human brain tumor-initiating cells (BTICs) with conditioned medium from primary human fetal neural progenitor cells (hfNPCs). We identified that treatment with hfNPC-conditioned medium significantly increased GBM BTIC viability over time compared to non-treated (NT) or BTIC-conditioned medium (**Supplementary Figure 1A**). For further exploration of the BTIC-hfNPC interaction, BTICs were co-cultured with hfNPCs using semi-permeable Transwell chamber inserts. Through Ki67 staining, we found that co-culture with hfNPCs increased the proliferation percentage of multiple BTIC lines compared to both non-treated (NT) lines and BTIC-BTIC co-culture controls (**Supplementary Figure 1B**). Additionally, hfNPCs induced a significant increase in BTIC migration compared to control conditions via Transwell migration assay (**Supplementary Figure 1C**). In the opposite conditions, co-culture with BTICs resulted in significantly decreased proliferation of hfNPCs compared to self-co-culture or NT groups (**Supplementary Figure 1D**). These results indicate that there is a direct interaction between the neurogenic cell population of the SVZ and GBM BTICs which contributes to increased tumor malignancy and decreased neurogenesis.

### Neural stem/progenitor cell-specific proteomics of the SVZ reveals decreased neuronal maturation in the presence of LV-proximal GBM

Due to the observed alterations in neurogenesis and NSC cell fate via IHC, we next wanted to examine changes in the NSC/NPC proteome dependent on GBM proximity to the LV using the MetRS L274G nascent protein labeling system. In Nes-MetRS* mice, we first ensured that our Cre driver was correctly inducing GFP expression in Nestin+ cells only following TAM administration. We injected Nes-MetRS* mice with TAM or corn oil intraperitoneal (IP) for five days and examined brain tissue for GFP expression 2 weeks later. No GFP+ cells were found within corn oil-injected animals, while Nestin+/GFP+ cells were identified at high levels above background within the SVZ upon TAM administration (**Supplementary Figure 2A**), as well as in the rostral migratory stream and sparsely in the dentate gyrus of the hippocampus (**Supplementary Figure 2B-C**). We next confirmed that the non-canonical methionine analog, azidonorleucine (ANL), was being incorporated into proteins in NSC/NPC-specific manner. Two weeks after TAM administration in Nes-MetRS* mice, ANL was provided IP over five days, followed by fluorescent non-canonical amino acid tagging (FUNCAT) analysis of the SVZ. Only ANL-injected animals had significant FUNCAT signal, which was mainly limited to GFP+ NPCs of the SVZ and the surrounding area (**Supplementary Figure 2D**). Additionally, via bio-orthogonal non-canonical amino acid tagging (BONCAT) we found that ANL-injected animals with the mutated MetRS enzyme had increased cell-specific metabolic protein labeling over background (**Supplementary Figure 2E**) and resulted in successful pulldown of higher levels of clicked, biotinylated proteins in animals induced with TAM and injected with ANL (**Supplementary Figure 2F**).

We then activated NSC labeling with TAM and injected animals intracranially with either LV-proximal vehicle, LV-distal GBM, or LV-proximal GBM. ANL was then delivered to animals and the ipsilateral SVZ was extracted for NSC/NPC cell-specific label-free quantitative (LFQ) proteomic analysis (**Figure 2A**). Through an analysis of five replicates per group, we identified a total of 2,669 NPC-specific proteins from the SVZ. After performing enrichment analysis with an ANOVA p-value cutoff of 0.05, there were 74 differentially expressed proteins (DEPs) in the SVZ of LV-proximal GBM animals compared to LV-proximal vehicle animals, and 84 DEPs in LV-proximal GBM SVZ compared to LV-distal GBM SVZ. Of these proteins, there were a total of 22 overlapping DEPs (18 downregulated and 4 upregulated) in LV-proximal GBM between the two analyses (**Figure 2B-C**). Of the 18 downregulated proteins, 12 are related to nervous system development, neuronal signaling, and synaptic function, such as Ache, Grm3, Grm5, Slc1a2 (EAAT2, glutamate transporter 1), and Slc17a7 (VGLUT1). STRING analysis and analysis of gene ontology (GO) terms and KEGG pathway enrichment were performed for the DEPs (**Figure 2D-E**). Commonly altered terms were largely related to transmembrane transport, synaptic transmission, neural cell projection, and calcium signaling (**Figure 2E**), all of which are important for the function of mature neurons^43^. This confirms histological findings of decreased neurogenesis and neuroblast differentiation in the presence of an LV-proximal GBM and implicates the identified shared altered proteins in regulating neuronal maturation of NPCs.

**Figure 2.**
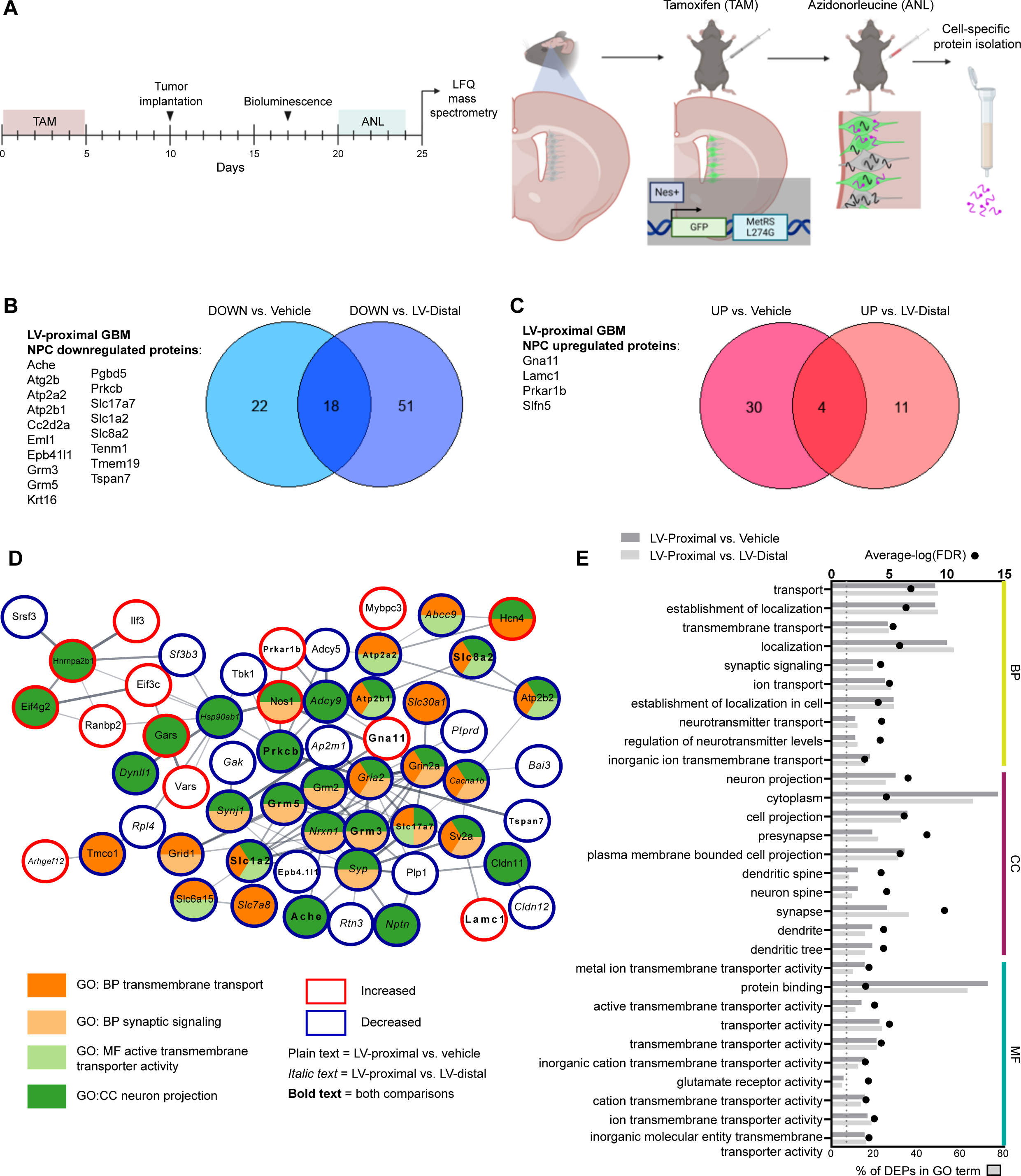
Cell-specific quantitative proteomics of SVZ NPCs reveals decreased expression of neuronal maturation proteins in the presence of LV-proximal glioma. (A) Schematic illustrating experimental timeline and methodology for Figure 2. (B) Venn diagram and list of downregulated proteins in the SVZ of LV-proximal GBM animals compared to vehicle and LV-distal GBM controls. (C) Venn diagram and list of upregulated proteins in the SVZ of LV-proximal GBM animals compared to vehicle and LV-distal GBM controls. (D) STRING interaction network for a subset of the LV-proximal GBM associated DEPs that belong to the indicated GO or KEGG terms or directly interact with proteins belonging to these terms. Node colors indicate protein affiliation with the indicated terms. Plain text indicates a DEP in the LV-proximal GBM vs. vehicle interaction. Italic text indicates a DEP in the LV-proximal GBM group vs. LV-distal GBM group. Bold text indicates the DEP overlaps between the two interactions. (E) Overlapping GO terms between the two interaction DEPs. BP = biological process, CC = cellular compartment, MF = molecular function.

### LV-proximal GBM induces DNA damage-mediated senescence in SVZ NPCs

One of the significantly increased proteins in the NPC proteome in response to LV-proximal GBM is schlafen family member 5 (Slfn5, **Figure 3A**). To confirm the increased expression of this protein in NSCs/NPCs in the presence of LV-proximal tumors, tissue from Nes-MetRS* mice was probed for Slfn5 expression. Slfn5 was found to be increased in NPCs in the presence of LV-proximal GBM at the tissue level via IHC (**Figure 3B-C**), confirming a cell-specific Slfn5 increase in NPCs and their progeny. Recently, Slfn5 has been identified to participate in reorganizing chromatin to facilitate non-homologous end joining of double strand breaks in DNA^44^. Slfn5+ puncta were confirmed to lie within nuclei of GFP+ cells of the SVZ (**Figure 3C**, inset), suggesting the presence of DNA damage and repair mechanisms at work in the SVZ of LV-proximal GBM-bearing animals. To examine whether LV-proximal GBM induces DNA breakage in the SVZ, we performed a TUNEL assay on the Nes-MetRS* brain tissue. We identified that compared to LV-distal GBM or LV-proximal vehicle controls, LV-proximal GBM induced a significant increase in the percent of TUNEL+ cells in the SVZ (**Figure 3D-E**), confirming an accumulation of DNA damage. The DNA damage response results in halted proliferation while promoting either apoptosis or cellular senescence^45^, depending on the severity of the damage and activation of downstream pathways. Our previous work identified no significant changes in apoptosis of SVZ cells in the presence of nearby GBM, despite decreased stem cell number and decreased proliferation^24^. We therefore decided to investigate alterations in senescence using the marker p21, a regulator of the DNA damage response in senescent cells^46^. In the Nes-MetRS* mouse model, we identified an increased percentage of p21+ SVZ nuclei in the presence of LV-proximal glioma (**Figure 3F-G**). Together, these data indicate that LV-proximal GBM induces increased DNA damage and senescence in NPCs of the SVZ.

**Figure 3.**
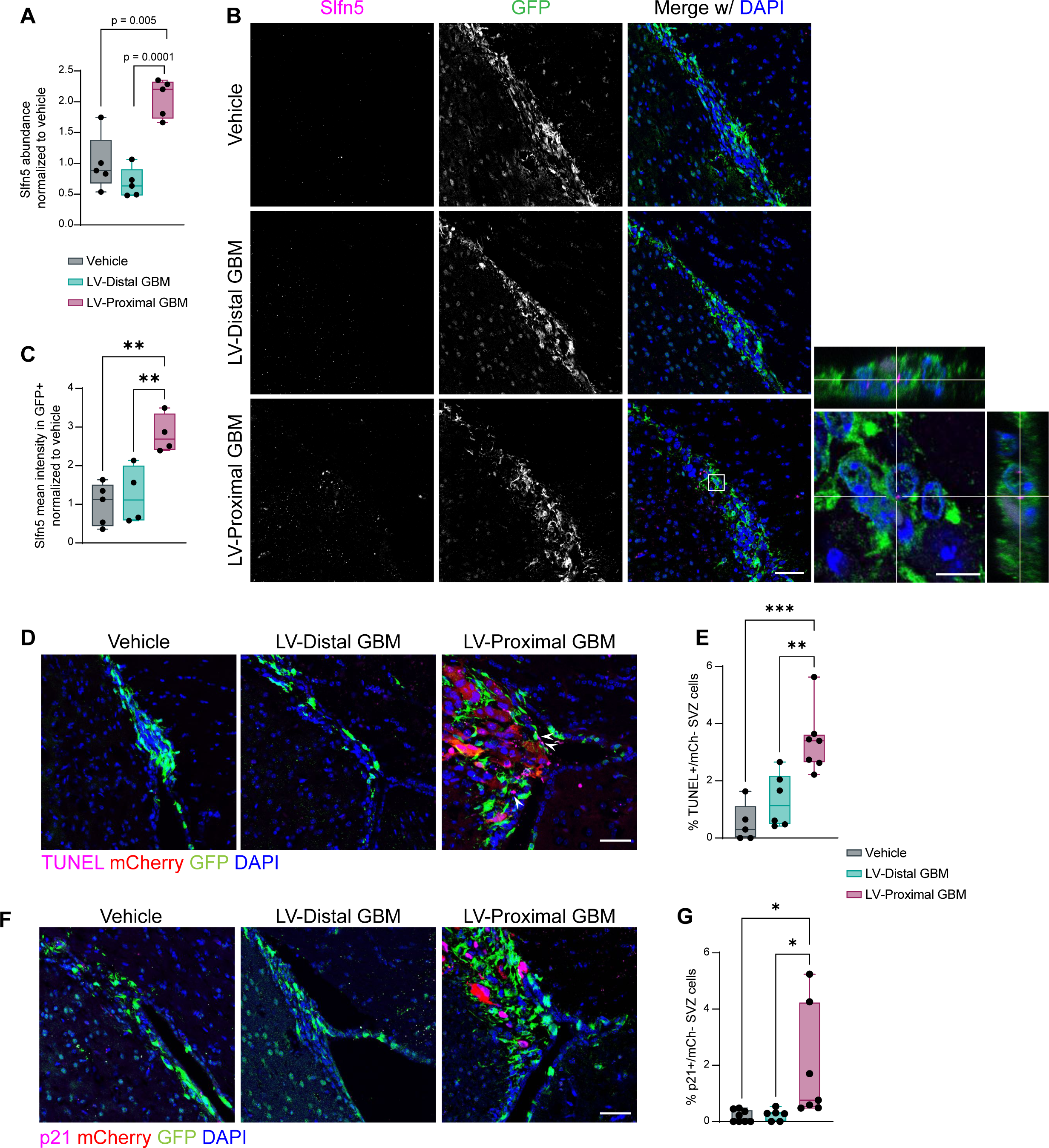
LV-proximal GBM induces increased DNA damage and senescence in SVZ NPCs. (A) LFQ proteomic results for Slfn5 quantification (n = 5). Normalized to vehicle group. (B) Representative immunofluorescent images of increased Slfn5+ in GFP+ cells of the LV-proximal GBM SVZ. Scale bar = 25 µm. White box indicates where inset on right is taken, showing Slfn5+ puncta within GFP+ cells. Scale bar for inset = 10 µm. (C) Slfn5+ mean intensity quantification in GFP+ cells. Normalized to vehicle group. (D) Representative immunofluorescence images of increased TUNEL+ cells in LV-proximal GBM SVZ. Scale bar = 50 µm. White arrows indicate TUNEL+/mCh-nuclei. (E) Quantification of percentage of TUNEL+ cells in SVZ across groups. (F) Representative immunofluorescence images of increased p21+ cells in LV-proximal GBM SVZ. Scale bar = 50 µm. (G) Quantification of percentage of p21+ cells in SVZ across groups. p < 0.05 *, p < 0.01 **, p < 0.001 ***, p < 0.0001 ****

### BTICs increase expression of pro-malignancy proteins, including cathepsin B, upon co-culture with NPCs

To analyze the direct influence of NPCs on GBM BTIC protein expression in the absence of other brain microenvironmental factors such as cerebrospinal fluid (CSF), we performed cell-specific proteomics of BTICs using the MetRS* labeling system in co-culture. Patient-derived BTICs were transduced with lentivirus encoding the mutant MetRS and co-cultured with wild type BTICs or hfNPCs (**Figure 4A**). LFQ proteomics of three independent replicates identified 9,452 peptides corresponding to 1,058 different BTIC-specific labeled proteins. Following enrichment analysis with a cutoff threshold of 1.5 fold-change, we identified 54 proteins downregulated and 52 proteins upregulated by NPC coculture (**Figure 4B**). GO and KEGG analysis of BTIC-specific DEPs revealed strong biological process signatures related to developmental biology and cell migration, including actin cytoskeleton organization, developmental process, actin filament-based process, and system development (**Figure 4C-D**). Additionally, several of the proteins are related to extracellular cellular compartment terms such as extracellular exosome, extracellular vesicle, and extracellular space (**Figure 4C-D**), implicating changes in intercellular signaling due to co-culture with NPCs. The DEP most increased by NPC co-culture and that can be both cell-contained and secreted extracellularly in GBM is the lysosomal cysteine protease cathepsin B (CTSB). We confirmed proteomic findings of increased CTSB in co-culture by performing western blot on BTIC-specific proteins (**Figure 4E-F**). We also identified increased expression of CTSB in LV-proximal GBM compared to LV-distal counterparts in our rodent model (**Figure 4G-H**). Together, these analyses identify several malignancy- and migration-related proteins upregulated in BTICs upon exposure to NPC and among these implicates CTSB as a potential target for LV-proximal GBM.

**Figure 4.**
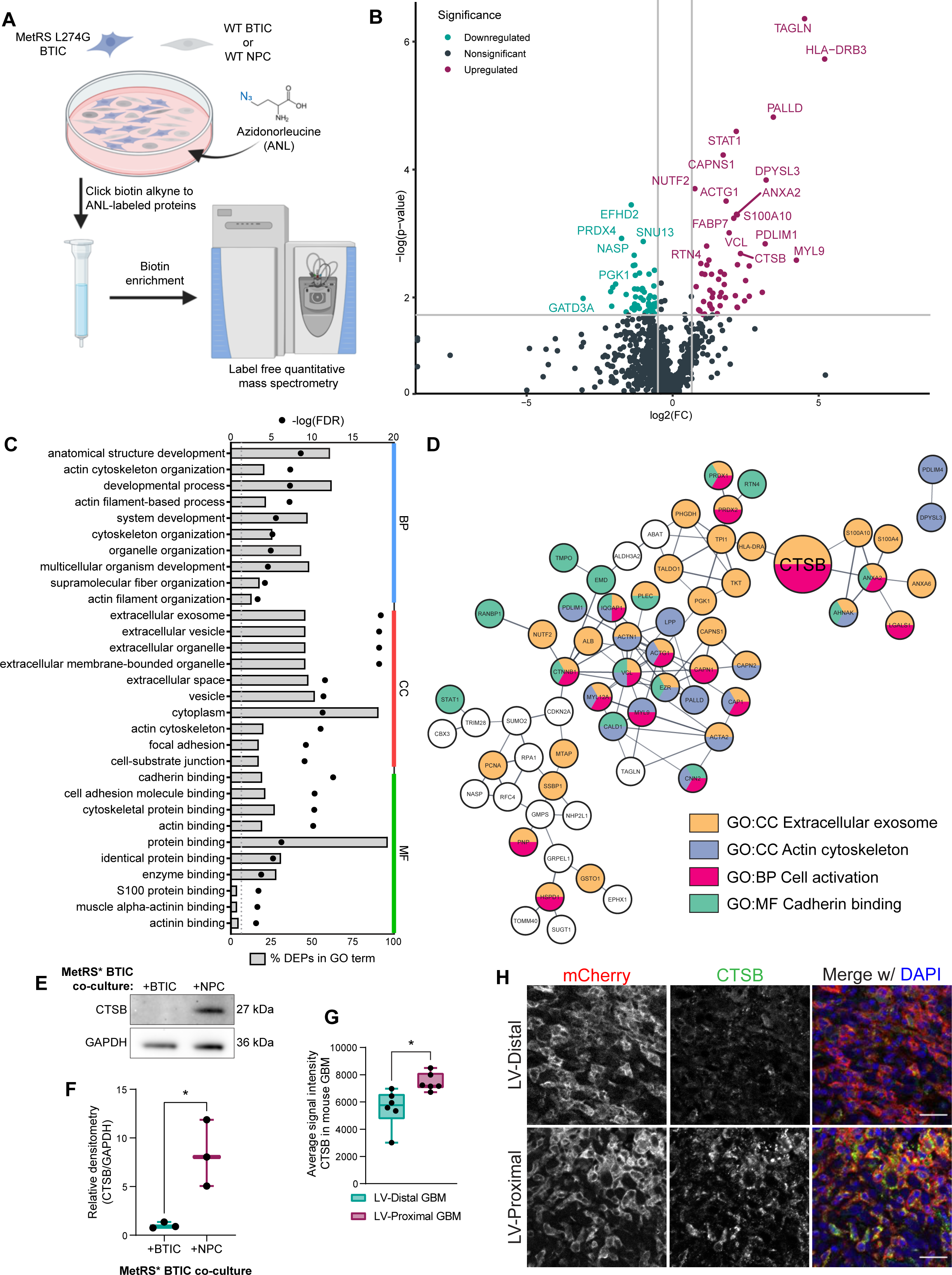
BTIC co-culture with hfNPCs results in GBM-specific increase in the expression of pro-malignancy proteins including cathepsin B. (A) Schematic illustrating co-culture setup. (B) Volcano plot of BTIC-specific proteins upregulated and downregulated by co-culture with NPCs. Gray lines indicate fold change = 1.5 (x-axis) and adjusted p-value at 0.05 (y-axis). (C) Top ten gene ontology terms in each GO category associated with DEPs from BTIC co-culture proteomics. Both the % of DEPs (bars) and false discovery rate (points) of terms are included. BP = biological process, CC = cellular compartment, MF = molecular function. (D) STRING interaction network for a subset of the NPC co-culture associated DEPs that belong to the indicated GO or KEGG terms or directly interact with proteins belonging to these terms. Node colors indicate protein affiliation with the indicated terms. CTSB highlighted by larger size. (E) Representative western blot for CTSB in BTIC-specific proteins from control co-culture and NPC co-culture. GAPDH is shown as a loading control. (F) Quantification of western blot densitometry for BTIC-specific CTSB normalized to GAPDH (n = 3). (G) Quantification of IHC mean intensity of CTSB within murine GBM (n = 6). (H) Representative IHC of mouse tumors showing increased CTSB in LV-proximal GBM. Scale bar = 25 µm p < 0.05 *, p < 0.01 **, p < 0.001 ***, p < 0.0001 ****

### Cell-intrinsic and soluble cathepsin B contribute to GBM malignancy

Due to the increase in CTSB expression in BTICs upon co-culture with NPCs, we next explored the effects of both cell-intrinsic and secreted CTSB on BTIC biology. To test the effect of endogenous CTSB on patient-derived BTICs, we performed lentiviral knockdown (KD) of CTSB in three BTIC lines using two short hairpin RNA (shRNA) constructs. An empty vector (EV) construct was also used as an experimental control. Following transduction, we confirmed CTSB KD using western blot (**Figure 5A**). We then investigated how CTSB intrinsic expression affects the malignant behavior of BTICs in culture. Knockdown of CTSB decreased viability (**Figure 5B**) and proliferation rate (**Figure 5C**) compared to EV controls, indicating that CTSB strongly contributes to the cellular growth of human GBM BTICs. Additionally, CTSB KDs exhibited decreased transwell migration (**Figure 5D**), total distance migrated, and distance traveled from origin (**Figure 5E-G**) in timelapse migration studies. In examining stem cell frequency using limiting dilution assay (LDA) to test for self-renewal capacity, we found that CTSB-silenced BTICs had reduced stem cell frequency compared to EV controls (**Figure 5H**). These results demonstrate that silencing the expression of endogenous CTSB reduces the stem cell fraction and the proliferation, viability, and migration capacity of GBM cells.

**Figure 5.**
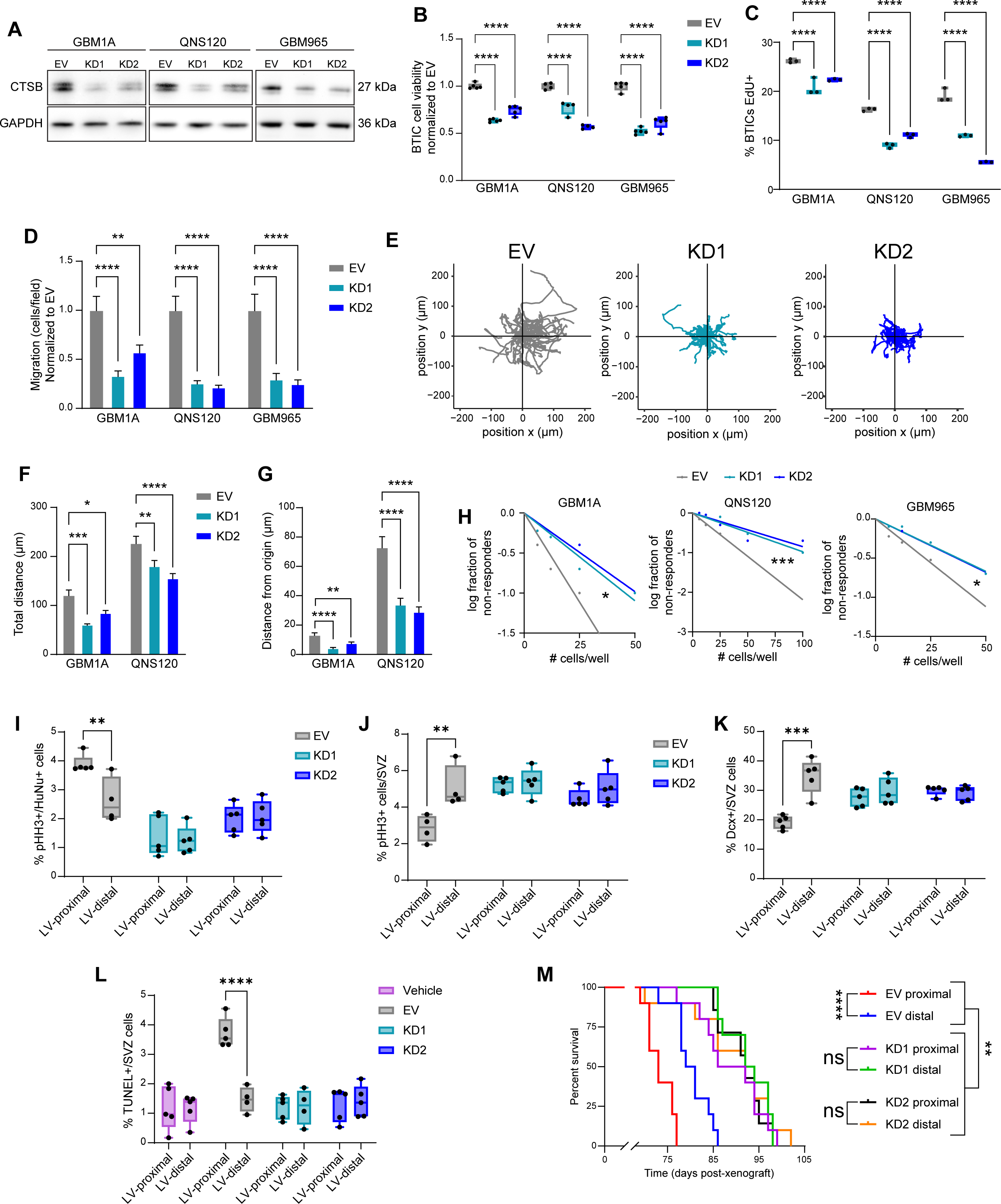
Cell-intrinsic CTSB contributes to GBM malignancy and LV-proximal GBM phenotypes. (A) Western blot confirming CTSB KD in BTIC lines. GAPDH is shown as a loading control. (B) Cell viability measurements for EV and CTSB KD at 48h after addition of alamarBlue reagent (n = 5). Normalized to EV. (C) Proliferation assay indicating percentage of EdU+ BTICs in EV and CTSB KD (n = 3). (D) Transwell migration of EV and CTSB KD. (E) Representative plots of migration from origin for EV and CTSB KD in BTIC line QNS120. (F) Total distance traveled by EV and CTSB KD in timelapse migration. (G) Distance from origin traveled by EV and CTSB KD in timelapse migration. (H) LDA self-renewal analysis for EV and CTSB KD. (I) Percentage of proliferation in different shRNA GBMs proximal or distal to LV (n = 4-5). (J) Percentage of proliferation in the SVZ in the presence of different shRNA GBMs proximal or distal to LV (n = 4-5). (K) Percentage of Dcx+ neuroblasts in the SVZ in the presence of different shRNA GBMs proximal or distal to LV (n = 5). (L) Percentage of TUNEL+ cells in the SVZ in the presence of PBS vehicle injection or different shRNA GBMs proximal or distal to LV (n = 4-5). (M) Kaplan-Meier curves of mice bearing different shRNA GBMs proximal or distal to LV (n = 7-10) p < 0.05 *, p < 0.01 **, p < 0.001 ***, p < 0.0001 ****

To examine how tumor-derived CTSB contributes to GBM prognosis, either the EV or CTSB KD BTIC lines were injected at LV-distal and LV-proximal locations in immunocompromised athymic nude mice. While EV control GBM tumors resulted in the previously observed increased tumor proliferation and decreased SVZ proliferation when injected in LV-proximal locations, CTSB KDs did not induce decreased SVZ proliferation and had significantly lower GBM cell proliferation irrespective of injection location (**Figure 5I-J**). CTSB KD in tumors also prevented the loss of Sox2+ stem cells in the SVZ and increase in Sox2+ tumor cells seen in control LV-proximal GBM (**Supplementary Figure 3A-B**). Additionally, while LV-proximal EV GBMs resulted in significantly decreased Dcx+ neuroblasts in the SVZ compared to LV-distal EV GBMs, CTSB KD tumors did not significantly alter neuroblast production (**Figure 5K**), implicating GBM-derived CTSB in decreased neuronal differentiation. SVZ cell DNA damage was also prevented by CTSB KD, where LV-proximal GBM no longer induced increased TUNEL+ cells in the SVZ (**Figure 5L**), suggesting GBM CTSB may play a role in mediating microenvironmental damage. Within the GBM, CTSB KD tumors had significantly increased TUNEL+/HuNu+ cells, showing increased DNA damage within the tumors themselves (**Supplementary Figure 3C**). Remarkably, the silencing of CTSB in BTICs prolonged median survival overall in comparison to EV controls, and also abolished the malignancy promoting effect of LV proximity (**Figure 5M**). These results indicate that cell-intrinsic CTSB plays a major role in GBM progression, and that the effect of the SVZ on GBM malignancy is mediated in large part by GBM CTSB.

To determine how soluble CTSB affects GBM BTIC biology, recombinant human CTSB was added to BTIC culture media. We determined that the addition of soluble CTSB increased both the viability (**Supplementary Figure 4A**) and proliferation rate (**Supplementary Figure 4B**) of BTICs. BTIC migration was also increased by soluble CTSB in transwell (**Supplementary Figure 4C**) and timelapse migration assays (**Supplementary Figure 4D-F**). These results show that secreted CTSB in the microenvironment also contributes to increased BTIC malignancy via autocrine and paracrine signaling.

### Soluble CTSB promotes NPC senescence

To evaluate how BTIC-secreted CTSB may contribute to the disruption in neurogenesis, we treated hfNPCs with recombinant CTSB in the culture medium. Interestingly, although cell viability increased in hfNPCs with the addition of soluble CTSB (**Figure 6A**), proliferation rate was decreased (**Figure 6B**), matching the observed *in vivo* phenotype. The migration activity of hfNPCs was also increased in the presence of soluble CTSB as observed in transwell (**Figure 6C**) and timelapse migration (**Figure 6D-F**), implicating a pro-migratory role of soluble CTSB on NPCs. Previous studies have identified increased CTSB expression and lysosome number in low-proliferating NSCs compared to high-proliferating progenitor cells^47^. Additionally, lysosomes are increased in size upon SVZ aging^47^, suggesting a potential role of CTSB and lysosomes in increased senescence. Therefore, we next tested whether soluble CTSB contributes to senescence or lysosomal phenotypes *in vitro*. Treatment with recombinant CTSB resulted in increased β-galactosidase expression in NPCs, identified via Senescence Green assay (**Figure 6G-H**). Additionally, we used LysoTracker to quantify the number and size of lysosomes in treated NPCs. Recombinant CTSB resulted in increased lysosome number and size in NPCs (**Figure 6G, I-J**). These results together suggest that soluble CTSB push NPCs toward a more senescent and less proliferative phenotype *in vitro*, strongly suggesting that GBM CTSB drives this phenotype in the SVZ of LV-proximal GBM bearing animals.

**Figure 6.**
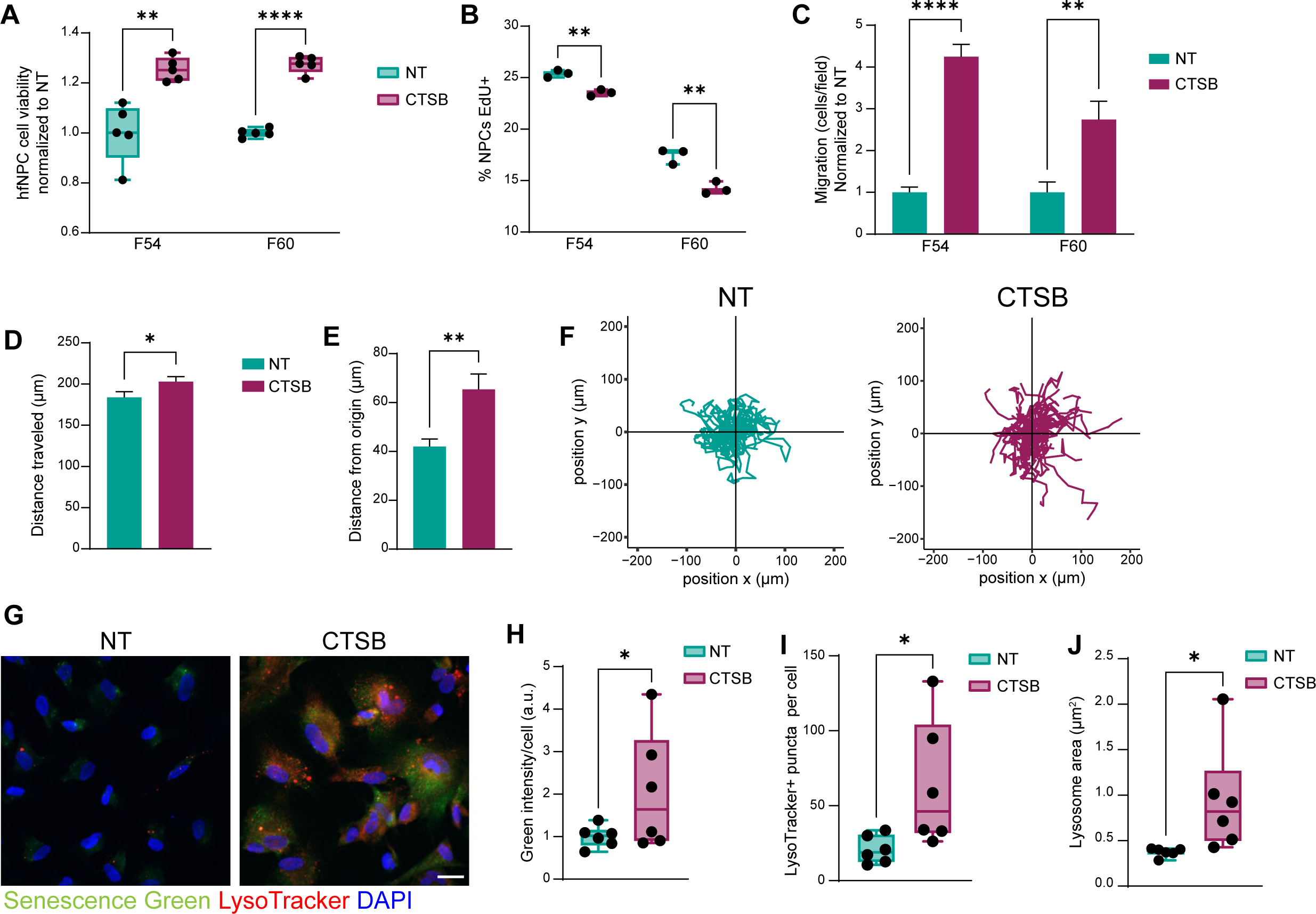
Soluble CTSB contributes to decreased proliferation and increased senescence of hfNPCs. (A) hfNPC cell viability measurements for NT and +CTSB at 48h (n = 5). (B) Proliferation assay indicating percentage of EdU+ hfNPCs in NT and +CTSB conditions (n = 3). (C) Transwell migration of hfNPCs NT and +CTSB. (D) Total distance traveled by hfNPCs F54 NT and +CTSB in timelapse migration. (E) Distance from origin traveled by hfNPCs F54 NT and +CTSB in timelapse migration. (F) Representative plots of hfNPC migration from origin when NT or +CTSB in line F54. (G) Representative immunofluorescent images of F60 with Senescence Green (β-galactosidase) and LysoTracker labeling in NT or +CTSB conditions. Scale bar = 25 µm. (H) Quantification of Senescence Green levels in F60 NT or +CTSB (n = 6). Normalized to NT. (I) Quantification of number of LysoTracker+ lysosomes per cell (n = 6). (J) Quantification of LysoTracker puncta area per well (n = 6). p < 0.05 *, p < 0.01 **, p < 0.001 ***, p < 0.0001 ****

### Patient role of CTSB in LV-contacting GBM

We have thus far shown that the interaction between BTICs and NPCs results in a malignancy-promoting upregulation of CTSB in culture and in our rodent model. However, due to the decreased frequency of neurogenesis in the adult human SVZ compared to rodents^11,12^, it is unclear whether these findings apply to human cases. To evaluate the contribution of CTSB to glioma patient outcome, we analyzed the TCGA GBM and LGG databases using the GlioVis data visualization tool^48^. We found that CTSB transcript is increased in GBM tissue compared to normal brain (**Figure 7A**). CTSB is also increased by glioma grade (**Figure 7B**) and is associated with decreased overall survival in the TCGA GBM cohort (**Figure 7C**) at the mRNA level. Additionally, we confirmed that CTSB protein is upregulated in GBM tissues compared to non-tumor cortex via western blot (**Figure 7D-E**). Finally, to determine whether CTSB protein is upregulated by proximity to the LV, we obtained intraoperative patient-matched surgical navigation-guided biopsies of GBM at locations close and far from the LV in patients with SVZ-contacting tumors (n = 8 patients) (**Figure 7F, Supplementary Table**). We found that CTSB protein is increased in LV-proximal GBM biopsies compared to the patient-matched LV-distal biopsies (**Figure 7G-H**), confirming that the LV or SVZ microenvironment upregulates CTSB protein expression in GBM patients.

**Figure 7.**
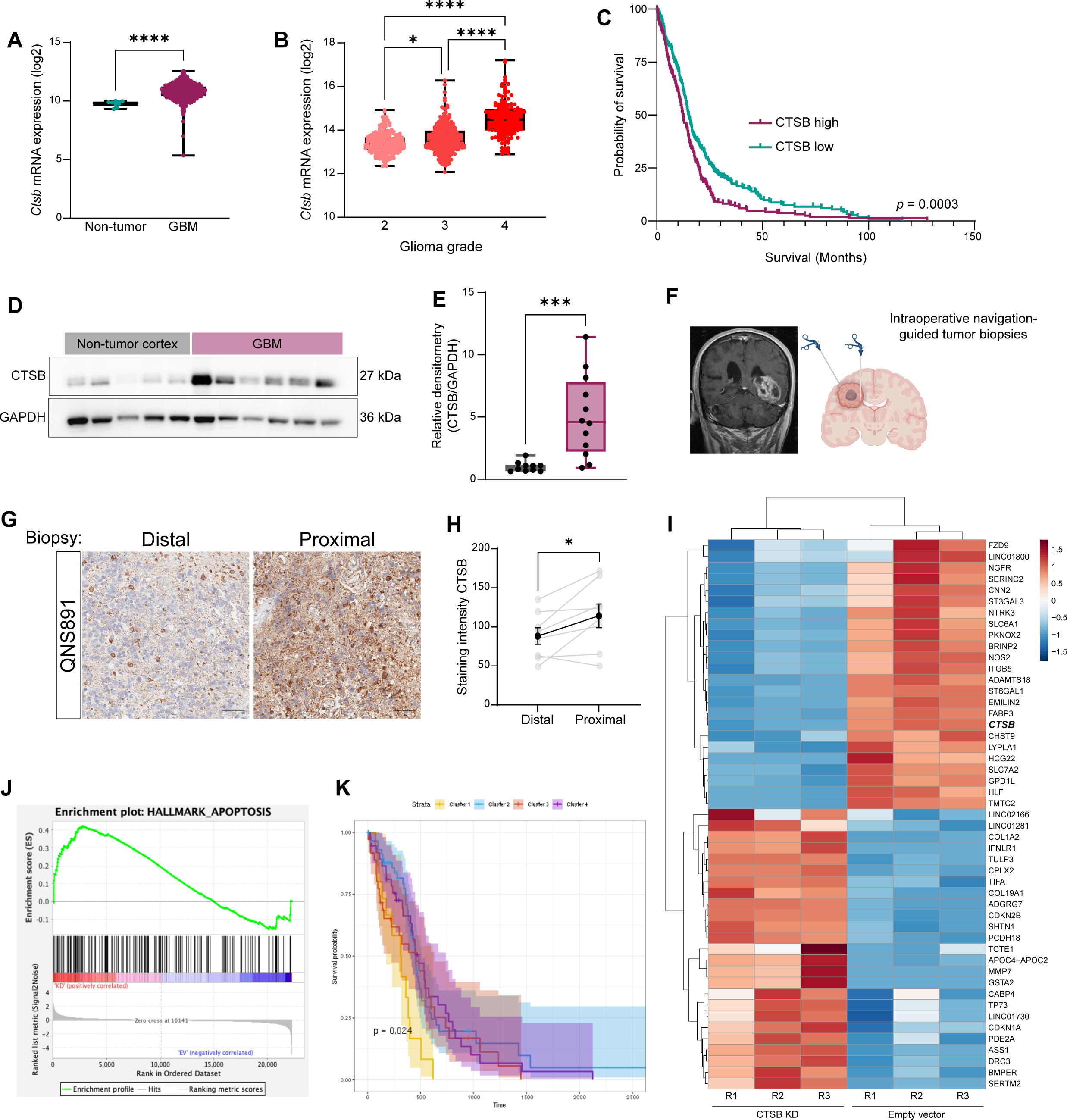
Role of CTSB in patient LV-contacting GBM. (A) CTSB gene expression in non-tumor and GBM tissues from TCGA database. (B) CTSB gene expression among different glioma grades from TCGA database. (C) Kaplan-Meier curves of survival outcomes for CTSB high and CTSB low GBM. Groups separated at median expression. (D) Representative western blot for CTSB in patient-derived non-tumor cortex and GBM tissues. (E) Quantification of western blot densitometry data from patient tissues (n = 10-12). Normalized to non-tumor cortex. (F) Schematic outlining collection of matched patient-derived biopsies from LV-contacting tumors. (G) Representative staining for CTSB in matched patient-derived LV-distal and LV-proximal biopsies for GBM patient QNS891. (H) Quantification of CSTB staining intensity in matched LV-distal and LV-proximal GBM biopsies (n = 8). (I) Gene heatmap of the 25 most downregulated and upregulated genes in CTSB KD compared to EV controls (n = 3). CTSB gene indicated in bold text. (J) GSEA plot for apoptosis hallmark gene set indicating increase in CTSB KD. (K) TCGA GBM patient Kaplan-Meier curves by cluster showing one cluster with significantly decreased median survival. p < 0.05 *, p < 0.01 **, p < 0.001 ***, p < 0.0001 ****

To determine how CTSB gene expression contributes to gene signatures, we performed bulk RNA sequencing on EV vs. CTSB KD1 GBM BTICs. A variety of genes were found to be regulated by CTSB expression in BTICs (**Figure 7I**), including several previously identified to contribute to BTIC biology. Upon performing gene set enrichment analysis (GSEA) for hallmark gene sets in the Broad Institute Molecular Signatures Database (MSigDB), we observed a strong upregulation of the apoptosis gene signature in CTSB KD compared to control (**Figure 7J**). We then used this gene signature to cluster TCGA GBM patients with RNA sequencing data by apoptosis signature gene expression (**Supplementary Figure 5**). Through this analysis, we identified cluster 1 to have significantly decreased survival compared to the other three clusters (**Figure 7K**).

## Discussion

The distinct pathobiology and specific molecular mechanisms driving LV-contacting GBM is of great clinical importance, due to the increased aggressiveness of these tumors resulting in worse patient outcomes. In this study, we determined that proximity of GBM to the SVZ contributes to both increased GBM malignancy and decreased neuronal differentiation in an immunocompetent rodent model. Through cell-specific proteomics, we determined that LV-proximal GBM reduced expression of several proteins related to neuronal maturation and synaptic function in SVZ NPCs while increasing DNA damage and senescence. On the GBM side, co-culture with NPCs results in cell-specific upregulation of CTSB. We have shown that cell intrinsic CTSB contributes to the malignant phenotype of GBM BTICs *in vitro* and *in vivo*, while soluble CTSB also plays a role in promoting BTIC proliferation and migration. Finally, we show that soluble CTSB may be one of the molecules stalling NPC differentiation via upregulating senescence, and that CTSB is also an important molecule in LV-contacting GBM in the patient population. Together, these data implicate the bidirectional interaction between NPCs and GBM as an important contributor to tumor malignancy and decreased brain health.

Interestingly, our data show that exogenous CTSB induces senescence in NPCs, shifting them to a less proliferative and more aged phenotype. Previous work has shown that the CTSB gene is upregulated in quiescent NSCs compared to activated transit amplifying cells and their progeny^47^. This is likely due to lysosomal CTSB; it was found that quiescent NSCs contained larger lysosomes with protein aggregates compared to activated NSCs, which were even further increased in size upon aging^47^. Indeed, the expression of lysosomal biosynthesis regulators contributes to maintaining the stemness of NPCs, where knockdown of these genes or lysosomal transporters results in premature neuronal differentiation^49^. When we treated NPCs with soluble CTSB, we observed an increase in the number and size of the lysosomes alongside increasing senescence. This is an interesting area for further study, as it is still unknown how exposure to soluble CTSB affects lysosomal dynamics in NPCs, or if soluble CTSB is internalized in these cells. Furthermore, it remains to be studied how senescence in NPCs may alter functional neurogenesis outcomes, such as neuroblast migration down the Rostral Migratory Stream (RMS) and neuronal maturation in the olfactory bulb. These areas are important areas of future study to fully understand the effect of CTSB on NPC biology.

Our data show that glioma tumor proximity to the SVZ results in the downregulation of several proteins involved in neuronal maturation and synaptic function. This is interesting, as it shows that the endogenous NPCs of the SVZ likely do not contribute to the newly discovered neuron-glioma synapses in the field of cancer neuroscience^50–53^. This could be due to two potential explanations; either these neuron-glioma synapses are less important in LV-proximal GBM, or these tumors receive neuronal input from other pre-existing neurons in the nearby striatum while maintaining a growth factor-rich niche in the presence of undifferentiated NSCs. Future work on the influence of cancer neuroscience on LV-proximal GBM will elucidate which of these hypotheses is true in these tumors.

Our work identifies CTSB as an important malignancy-promoting factor in LV-proximal GBM and that it is increased in tumor cells upon exposure to NPCs. It currently remains unknown what factor from NPCs induces the upregulation of CTSB in GBM cells, and whether this is dependent on cell-cell contact or is modulated by a secreted factor. Furthermore, the downstream effect of soluble CTSB on NPCs and the pathway through which it acts needs further exploration. The identification of the NPC-derived factor and how CTSB regulates NPC biological pathways need additional studies to fully understand.

Previously, CTSB expression in GBM cell lines has been closely linked to tumorigenic potential, invasiveness, and radioresistance^54–64^. These studies, however, have largely been performed using commercial cell lines and do not explore these findings in the context of patient heterogeneity. Here, we confirm the importance of CTSB in several low passage patient-derived BTICs and show that the cell-intrinsic expression of this molecule is important in the proliferation, invasiveness, and stemness properties of these cells. The contribution of CTSB to radioresistance in patient-derived lines should be further explored in future studies, as this would make this molecule an excellent target for new therapeutics. In addition to the cell-intrinsic contribution, we found that soluble CTSB also contributes to GBM cell growth and migration, suggesting additional malignancy-promoting properties of autocrine and paracrine signaling. CTSB can be released into the tumor microenvironment by a variety of cell types including endothelial cells, fibroblasts, and tumor-associated macrophages and myeloid-derived suppressor cells^65–68^. Additional cell-specific proteomic studies on other cell types of the tumor microenvironment will be essential for elucidating the role of soluble CTSB, as well as other molecules, on GBM progression.

Though this study focuses on the crosstalk between neoplastic cells and their microenvironment, there is no perfect *in vitro* or *in vivo* model to study this aspect of GBM. Therefore, we utilize several systems to confirm our findings. Our *in vitro* studies utilize co-culture between human BTICs and hfNPCs, but the interaction between these two cell types may be further mediated by a third cell type such as tumor-associated macrophages, or a non-cellular source of factors such as the CSF. In our transgenic model for MetRS* studies, we utilize an immunocompetent rodent with a syngeneic murine glioma line. Although Gl261 recapitulates several aspects of human GBM^69^, it is a commercial cell line and lacks the classic heterogeneity that is characteristic of the human disease. Therefore, in our knockdown experiments we implant patient-derived BTICs in immunosuppressed mice. This allows for a more heterogenous human glioma model, but these animals lack the functional immune system that further contributes to disease progression^26,70–76^. Currently, studying the tumor microenvironment in GBM necessitates the use of multiple models and crosschecks, as well as including patient samples to verify implicated pathways. Here we have combined the use of *in vitro* culture systems, immunodeficient and immunocompetent murine models, and patient samples to fully implicate the CTSB and senescence pathways in LV-proximal GBM. Hopefully with the technological and methodological growth of the field, such as the development of organoid co-culture^77–81^ and 3D biomimetic culture systems^82–84^, the simulated interaction of tumor cells with their microenvironment will more closely model the human condition.

Here we use a nascent proteomic labeling system, MetRS L274G in the presence of ANL, to explore the proteomic contributions in the crosstalk between GBM and NPCs. Although this system has been used in other areas of neuroscience^85–87^, cancer biology^88^, and stem cell biology^89–91^, we are the first to apply it in the context of neurogenesis or brain tumors. Our newly developed approach to this system can now be utilized to profile GBM proteomic dynamics *in vivo* and can be more widely applied to other diseases and microenvironmental niches. For example, NSCs play a role in other disorders such as neurodegenerative disease and stroke^92–94^, while GBM makes use of other malignancy-promoting niches such as the perivascular niche^95–97^. Proteomic characterization in these cell populations is extremely important due to post-transcriptional regulation of stem cell fate and low transcript-protein correlation^37,38,98^, meaning protein-based signaling analysis will give a more accurate picture of the biological pathways at play. Using the MetRS L274G system or other cell-specific proteomic isolation methods to further illuminate the proteomic dynamics of GBM-niche crosstalk and SVZ neurogenesis will lead to novel discoveries in the field and, potentially, the development of new therapeutic strategies.

Based on our results, we propose that NSC/NPCs contribute to GBM progression. This has some distressing implications for current clinical trials utilizing human NSCs for therapeutic delivery in GBM patients. Due to the homing nature of NSCs to tumor masses in vivo, several preclinical and clinical studies have used NSCs in stem cell therapies as delivery systems for oncolytic virus or as mediators of enzyme/prodrug therapies^99–102^. The clinical trials derived from these studies are still in early phases, though phase I results suggest no safety concerns with NSC-based treatment. If these treatments are successful, careful screening of individual NSC lines will need to be performed to ensure that each line used is not secreting large amounts of stemness-promoting growth factors, and that these lines do not result in CTSB upregulation in the patient glioma, which would then lead to downstream resistance to standard of care therapy.

Ultimately, our results indicate that there is malignancy-promoting crosstalk between GBM and the SVZ that disrupts normal neurogenesis through activation of senescence while promoting tumor progression through CTSB upregulation. Targeting this pathway using CTSB inhibitors may be a useful therapeutic approach to treat patients with these tumors. The future use of nascent proteomic labeling methods to perform cell-specific research in the tumor microenvironment will give significant insight into the biology of GBM.

## Supporting information

Supplemental table Patient demographics

## Acknowledgements

ESN was supported by NINDS F31NS120605, the Mayo Clinic Regenerative Sciences Training Program, and the Uihlein family scholarship award. ESN, LAW, VKJ, MNR, NZ, AC, and HGC were supported by NINDS K01NS110930 and the Uncle Kory Foundation. AQH was supported by the Mayo Clinic Clinician Investigator award, the William J. and Charles H. Mayo Named Professorship, the Monica Flynn Jacoby Endowed Chair, and the Uihlein Neuro-Oncology Research Fund. This publication was made possible by support of the Mayo Clinic Center for Biomedical Discovery (CBD), Mayo Clinic, Rochester, MN. We thank Brandy Edenfield for assistance in immunohistochemistry and scientific illustrator Stephen Graepel for creating the graphical abstract. Additionally, we thank Florine Collin and Jean Kanyo at the Keck MS & Proteomics Resource for support with proteomics sample preparation and data collection. We thank the MS & Proteomics Resource at Yale School of Medicine for providing the necessary mass spectrometers and the accompanying biotechnology tools funded in part by the Yale School of Medicine and by the Office of The Director, National Institutes of Health (S10OD02365101A1, S10OD019967, and S10OD018034). The funders had no role in study design, data collection and analysis, decision to publish, or preparation of the manuscript. ESN and MNR would like to thank the Mayo Clinic Graduate School of Biomedical Sciences for support.

## Author contributions

Conceptualization: ESN & HGC

Methodology: ESN, MMB, NZ, HGC

Resources: ESN, MMB, ARF, PS, WR, KLC, TTL, PZA, AQH, & HGC

Investigation: ESN, LAW, VKJ, MMB, MNR

Validation: ESN, LAW, VKJ, MMB, MNR

Formal analysis: ESN, LAW, MNR, DM, EJ, AC, YWA, TTL, HGC

Visualization: ESN, VKJ, DM, EJ

Project administration: ESN & HGC

Supervision: ESN, PS, WOR, PZA, AQH, HGC

Funding acquisition: ESN, PS, PZA, AQH, & HGC

Writing – original draft: ESN & HGC

Writing – review & editing: all authors

## Methods

### Experimental Animals

Animal experiments were fully approved by the Mayo Clinic Institutional Animal Care and Use Committee (protocols A00002260-16 and A00004969-19). Mice were housed in an AAALAC-accredited facility abiding by all federal and local regulations. For NSC proteomic labeling experiments, C57BL6/J Nestin-Cre/ERT2 transgenic mice (Nes-cre/ERT2, Jackson Laboratory, #016261)^103–105^ were crossed with STOPflox R26-MetRS L274G transgenic mice (STOPflox R26-MetRS*, Jackson Laboratory #028071)^106^. Animals were backcrossed in a C57BL6 background. Genotypes used for all NSC labeling experiments were Nestin^CreERT2/+^; MetRS^L274G/L274G^.

Cre activity for NSC proteomic labeling was induced via tamoxifen (TAM, Sigma-Aldrich #T5648). TAM dissolved in sterile corn oil was administered IP at a dosage of 140 mg TAM/kg body weight/day over a period of five days. Animals were monitored daily until the end of study for changes in weight and physical signs of toxicity or distress.

### Cell culture

All cell culture was performed in sterile conditions using aseptic technique. Cells were monitored for mycoplasma contamination once per month and were discarded if positive. Syngeneic murine glioma line 261 (GL261) was cultured in adherent conditions in DMEM media (Gibco) supplemented with 200 µM L-glutamine (GlutaMAX, Gibco), antibiotic-antimycotic solution, and 10% FBS. Patient-derived brain tumor initiating cells (BTICs) were cultured in complete media consisting of DMEM/F12 (Gibco) supplemented with antibiotic-antimycotic solution, B-27, 20 ng/mL epidermal growth factor (EGF; Peprotech), and 20 ng/mL fibroblast growth factor (FGF; Peprotech). Human fetal neural progenitor cells (hfNPCs) were cultured in complete media of DMEM/F12 supplemented with antibiotic-antimycotic solution, B-27, EGF, FGF, 10 ng/mL leukemia inhibitory factor (Millipore), and 5 µg/mL heparin. BTICs and hfNPCs were cultured as neurospheres in suspension until needed for experimentation. For adherent experiments, tissue culture flasks were coated with laminin (10 µg/mL in PBS) for 2h at 37C before plating cells.

### Intracranial tumor implantation

For NSC proteomic labeling experiments in mice of BL6 background, GL261 cells were transduced via lentivirus to stably express CMV-mCherry and CMV-luciferase PGK-puromycin^107^ (mCh-luc). pLV-mCherry was a gift from Pantelis Tsoulfas (Addgene plasmid #36084; http://n2t.net/addgene:36084; RRID: Addgene_36084). pLenti CMV Puro LUC (w168-1) was a gift from Eric Campeau & Paul Kaufman (Addgene plasmid #17477; http://n2t.net/addgene:17477; RRID: Addgene_17477). For human BTIC injection into athymic nude mice, cells were transduced to express PGK-GFP-IRES-LUC-W (GFP-luc)^108^. pHAGE PGK-GFP-IRES-LUC-W was a gift from Darrell Kotton (Addgene plasmid #46793; http://n2t.net/addgene:46793; RRID: Addgene_46793). Following transduction, intracranial implantation of cells was performed. Briefly, mice were placed under anesthesia and fitted into a stereotactic frame. For murine glioma experiments, 5.0×10^4^ GL261 mCh-luc+ cells in a volume of 1 μL PBS were injected into the right brain hemisphere. For human BTIC experiments, 3.5×10^5^ GFP-luc+ BTICs were implanted in 2 µl of PBS. Injection in LV-proximal and LV-distal locations was performed as previously reported^24^, but with a slightly more lateral LV-proximal injection. In mm relative to bregma, LV-proximal injections were AP: 1.0, L:1.35, and D: 2.3, while LV-distal locations were AP: 1.0, L: 2.1, and D: 2.3. Vehicle-injected mice (PBS only) at the LV-proximal location were included to account for NSC response to the intracranial injection. Tumor engraftment was monitored with bioluminescence following IP injection of luciferin. For lysate preparation, mice were intracardially perfused with 50 mL of ice cold 0.9% saline with 0.5 mg/mL heparin. For immunohistochemistry (IHC), animals were perfused with saline with heparin, then 4% paraformaldehyde (PFA) in PBS at study endpoint. For survival experiments, mice were maintained following GBM xenograft until reaching humane endpoint criteria defined as weight loss greater than or equal to 20% of body weight, inability to ambulate, inability to reach food or water, ataxia, paraplegia, inability to right oneself, or a body condition score of 1 or less using the IACUC approved scoring system.

### Azidonorleucine (ANL) administration

For animal experiments, azidonorleucine (ANL, Iris Biotech) was fully dissolved in sterile 0.9% saline solution. NaOH was added to pH = 7.4 and the ANL solution was filtered through a 0.22 µm pore syringe filter. ANL was administered IP to mice at a dosage of 200 mg/kg/d over five days. Non-labeled controls for NSC labeling experiments consisted of littermates that received only corn oil vehicle and ANL. For cell culture experiments, ANL was prepared in sterile DMEM/F12 to a 4 mM concentration of pH = 7.0.

### Immunohistochemistry (IHC)

For IHC analysis, paraffin-embedded brain sections (10 μm) were deparaffinized and rehydrated followed by antigen retrieval with sodium citrate buffer (10 mM, pH = 6.0). For fluorescent IHC, rehydrated sections or thawed frozen sections were washed, permeabilized, blocked with 10% normal goat serum, and incubated with appropriate primary antibodies at the dilutions described in Table 1 overnight at 4°C. The next day sections were washed and incubated in secondary antibodies diluted in blocking solution (1:500) for 1 hour at room temperature. Sections were counterstained with DAPI before coverslipping. For ICC, cells were washed twice, fixed with 4% PFA for 15 minutes, then went through the same staining protocol beginning at the permeabilization step. Preparations were imaged using a LSM 880 confocal microscope (Zeiss).

**Table 1.**

| Target | Manufacturer and catalog number | Host species | Use | Dilution |
| --- | --- | --- | --- | --- |
| Cathepsin B (CTSB) | Cell Signaling #31718S | Rabbit | IHC and Western | 1:100 – 1:1000 |
| Doublecortin (Dcx) | Cell Signaling #4604S | Rabbit | IHC | 1:500 |
| Glyceraldehyde-3-phosphate dehydrogenase (GAPDH) | Santa Cruz Biotechnology #sc-47724 | Mouse | Western | 1:5000 |
| Green fluorescent protein (GFP) | Aves Labs #GFP1010<br>ThermoFisher Scientific #A-11122 | Chicken<br>Rabbit | IHC<br>IHC | 1:500<br>1:500 – 1:1000 |
| Ki67 | ThermoFisher Scientific #MA5-14520 | Rabbit | IHC | 1:150 |
| Mash1 | BD Pharminigen #556604 | Mouse | IHC | 1:100 |
| Nestin | BD Pharminigen #556309 | Mouse | IHC | 1:100 |
| Olig2 | Millipore Sigma #AB9610 | Rabbit | IHC | 1:500 |
| Phospho-histone H3 (pHH3) | Cell Signaling Technology #9701S | Rabbit | IHC | 1:250 |
| S100β | Abcam #ab52642 | Rabbit | IHC | 1:100 |
| Slfn5 | Atlas Antibodies #HPA017760 | Rabbit | IHC | 1:100 |

### Cell viability assay

Cells were plated in a black 96 well plate with clear bottom and allowed to attach in complete media overnight. The following day, media was exchanged to base medium without growth factors. AlamarBlue viability reagent (Invitrogen) was added to total 10% of well volume. Fluorescence was measured with excitation/emission spectra 540/600. Viability was measured at 0, 4, 24, 48, and 72 hours following addition of reagent.

### Transwell chamber co-culture

Transwell chamber inserts with 0.4 μm pores (Corning) in either a 6 well plate or 24 well plate format were used to co-culture BTICs with hfNPCs. Cells were first allowed to adhere to the bottom of cell culture plates overnight in complete media. The next day, media was replaced with base media lacking growth factors and the Transwell inserts were placed on top. Cells to be co-cultured were placed into the top chamber in base media. Cells were incubated together for 48 hours before appropriate analysis.

### Transwell migration

Transwell chamber inserts with 8.0 μm pores (Corning) in a 24 well plate format were used to analyze migration. Based on optimization experiments, 4.0×10^4^ cells were placed into the top chamber. To ensure migration, an FBS gradient of 2.5% in the bottom chamber (500 μL) and 0.5% in the top chamber (250 μL) was established in media in the absence of added growth factors. Plates were incubated for 24 hours before processing. Cells remaining in top chamber were removed with a cotton swab, while cells that successfully migrated to the bottom of the membrane were fixed and labeled with DAPI before being imaged at 9 fields per membrane at a 10x magnification using a fluorescence microscope. Cells per field were quantified using ImageJ software.

### Fluorescent non-canonical amino acid tagging (FUNCAT) of newly synthesized proteins *in vivo*

FUNCAT labeling in tissue sections was performed according to previously published protocols^109^. Briefly, slides were washed and blocked as described for IHC. ANL-labeled proteins were clicked by mixing 200 μM triazole ligand (TBTA), 500 μM TCEP, 5 μM fluorescent alkyne tag, and 200 μM CuSO_4_ in PBS and adding to sections for 3.5 h at room temperature under gentle rotation. Slides were washed twice with PBS containing 1% Tween-20 and 500 μM EDTA for 15 minutes, followed by two washes with PBS containing 0.1% Triton. Slides were then counterstained using DAPI or underwent IHC staining beginning from the blocking step.

### Lysate preparation

For NSC MetRS* experiments, the ipsilateral SVZ to intracranial injection site was dissected out of the brain^110^ and wet tissue weight was measured. Tubes containing tissue were snap frozen in dry ice and stored at −80°C until lysate extraction. Tissue was manually homogenized in 12x volume of lysis buffer consisting of PBS with 1% SDS, 1% Triton X-100, 1x protease inhibitors, and Benzonase (≥250 U/mL). Lysates were sonicated and incubated at room temperature for 30 minutes to allow for complete lysis before centrifuging at 16,000 x g and retaining supernatant for downstream analysis.

For cell culture experiments, lysis buffer was added to flasks containing cells, which were then scraped and placed into tubes on ice. Lysates were sonicated and incubated on ice for 15 minutes with intermittent vortexing before centrifuging at 16,000 x g and retaining supernatant as whole cell lysate.

### Bio-orthogonal non-canonical amino acid tagging (BONCAT) and purification of cell-specific nascent proteome

For gel imaging, 100 μg of tissue lysate was first reduced with 25 mM dithiothreitol (DTT) at 80°C for 15 minutes. Following reduction, lysates were alkylated with 50 mM iodoacetamide (IAA, ThermoFisher) at room temperature for 1 hour and 30 minutes rotating at 1000 rpm and protected from light. This alkylation step was repeated for a total of two times. Proteins were then clicked to 10 μM TAMRA-DBCO using strain-promoted click chemistry for 1 hour at room temperature protected from light. Excess DBCO reagent was removed using Amicon Ultra Centrifugal Columns with 3 kDa cutoff (Millipore). A volume of 30 μL of resulting solution was run on an SDS-PAGE gel and imaged using the ChemiDoc MP System (BioRad). Imaging was followed by Coomassie Blue staining for total protein quantification.

For purification of ANL-labeled proteins, 1 mg of proteins were first reduced and alkylated as described for gel imaging. Proteins were then precipitated in ice-cold acetone and redissolved in fresh lysate buffer. Following protein concentration quantification by BCA, lysates were clicked to 10 μM DBCO-S-S-PEG3-biotin (BroadPharm) for 6 hours at room temperature. Excess DBCO reagent was removed with PD G-25 SpinTrap columns (Cytiva) and proteins were incubated with Streptavidin Sepharose High Performance affinity resin (Cytiva) for 16 hours at 4°C in rotation. Resin was washed three times with 1% Triton X-100 0.15% SDS in 1X PBS, three times with 1% Triton X-100 0.2% SDS in 1X PBS, and three times with 1X PBS. Proteins were eluted from beads using 50 mM DTT and 0.1% SDS in 1X PBS over 4 hours at room temperature.

### Label free quantitative (LFQ) mass spectrometry sample preparation

LFQ sample preparation was carried out similar to that previously described^111^, but with slight updated modification. The submitted samples were brought up to 100 µL with water. The proteins were precipitated utilizing an acetone precipitation procedure. Protein pellet was air dried, dissolved in 10 µL of 8M urea/0.4M ammonium bicarbonate, reduced with 1 µL of 45 mM dithiothreitol at 37°C for 30 minutes, and subsequently alkylated with 1 µL of 100 mM iodoacetamide at room temperature for 30 minutes in the dark. The solution was then diluted with 27 µL water and digested with 1 µL of 0.5 µg/µL trypsin at 37°C overnight. The tryptic digestion was quenched by acidifying with 2 µL 20% trifluoro-acetic acid (TFA). The peptide solution was then desalted using a mini Reverse Phase (RP) C18 desalting column (The Nest Group, Ipwich, MA). The eluted peptides were then dried in a SpeedVac and stored at −80°C. The dried samples were then re-dissolved in 5 μL 70% formic acid (FA) and 35 μL 0.1% TFA. An aliquot was taken to obtain total digested protein amount. A 1:10 dilution of Pierce Retention Time Calibration Mixture (Cat# 88321, ThermoFisher, Waltham, MA) was added to each sample prior to injecting onto the UPLC coupled Q-Exactive Plus mass spectrometer system for normalization of the LFQ data.

### LFQ data collection

Data dependent acquisition (DDA) liquid chromatography with tandem mass spectrometry (LC MS/MS) data collection for LFQ was performed on a Thermo Scientific Q-Exactive Plus mass spectrometer connected to a Waters nanoACQUITY UPLC system equipped with a Waters Symmetry® C18 180 μm × 20 mm trap column and a 1.7 μm, 75 μm × 250 mm nanoACQUITY UPLC column (35°C). The digests were diluted to 0.05 μg/μL with 0.1% TFA prior to injecting 5 μL of each triplicate analysis in block randomized order. To ensure a high level of identification and quantitation integrity, a resolution of 120,000 and 30,000 was utilized for MS and MS/MS data collection, respectively. MS and MS/MS (from Higher-energy C-Trap Dissociation (HCD)) spectra were acquired using a three second cycle time with Dynamic Exclusion on. All MS (Profile) and MS/MS (centroid) peaks were detected in the Orbitrap. Trapping was carried out for 3 min at 5 μL/min in 99% Buffer A (0.1% FA in water) and 1% Buffer B [(0.075% FA in acetonitrile (ACN)] prior to eluting with linear gradients that reach 25% B at 150 min, 50% B at 170 min, and 85% B at 175 min; then back down to 3% at 182 min. Two blanks (1st 100% ACN, 2nd Buffer A) followed each injection to ensure there was no sample carry over.

### Proteomic data analysis

The collected LC MS/MS LFQ data was processed with Progenesis QI software (Nonlinear Dynamics, version.4.2) with protein identification carried out using the Mascot search algorithm (Matrix Science, v. 2.7). The Progenesis QI software performs feature/peptide extraction, chromatographic/spectral alignment (one run was chosen as a reference for alignment), data filtering, and quantitation of peptides and proteins. A normalization factor for each run was calculated to account for differences in sample load between injections as well as differences in ionization. The normalization factor was determined by comparing the abundance of the spike in Pierce Retention Time Calibration mixture among all the samples. The experimental design was set up to group multiple injections from each run. The algorithm then tabulated raw and normalized abundances, and maximum fold change for each feature in the data set. The combined MS/MS spectra were exported as .mgf files (Mascot generic files) for database searching. The Mascot search results were exported as .xml files using a significance peptide cutoff of p < 0.05 (must have at least peptide count of "2", unique peptides count of "1", and Confidence score of greater than or equal to 35 (∼95% confidence in ID)) and protein FDR of 1%. The .xml search result and then imported into the Progenesis QI software, where search hits were assigned to corresponding aligned spectral features. Relative protein-level fold changes were calculated from the sum of all unique and non-conflicting, normalized peptide ion abundances for each protein on each run. Three to five biological replicates were processed for each condition.

Proteins that had a fold change ANOVA p-value < 0.05 were considered differentially expressed and were analyzed for enrichment of gene ontology (GO) terms and Kyoto encyclopedia of genes and genomes (KEGG) pathways using the g:Profiler web-based tool. For significance thresholds, the statistical domain scope was set to “Only annotated genes” and the Benjamini-Hochberg corrected FDR < 0.05. Protein-protein interaction networks were obtained using the STRING database^112^ with an interaction combined cutoff score of 0.7 and imported into the Cytoscape software platform^113^ for visualization.

### TUNEL assay

Fixed brain sections were rehydrated and underwent antigen retrieval before being processed for TUNEL assay using the Click-iT™ Plus TUNEL Alexa Fluor™ 647 Assay Kit for In Situ Detection (Invitrogen) according to manufacturer’s instructions. Unstained sections were included as a control.

### Immunoblotting

A total of 10-20 μg of proteins were subjected to electrophoresis on polyacrylamide gels and transferred to PVDF membranes. Blots were blocked in 5% BSA in TBST for 1 hour at room temperature before being incubated with the appropriate primary antibody in blocking buffer overnight at 4°C (see antibody table above). The following day, membranes were washed with TBST before being incubated with HRP-conjugated secondary antibody (1:5000 dilution) in blocking buffer for 1 hour at room temperature. Blots were then washed again with TBST, incubated with ECL substrate, and imaged using the MyECL Imager system (ThermoFisher) with CCD camera. Densitometry analysis was performed using ImageJ Fiji^114^.

### Lentiviral transduction

BTICs were adhered and transduced with lentiviral particles in the presence of polybrene (8 μg/mL) using an MOI of 100. Following infection, cells were selected for lentiviral incorporation using puromycin (0.5 μg/mL) over a period of at least six days. Knockdowns were confirmed before performing experimentation. All knockdown experiments were performed within four passages of lentiviral transduction.

### EdU proliferation assay

Proliferation of cells was measured using the Click-iT™ EdU Alexa Fluor™ 647 Flow Cytometry Assay Kit (Invitrogen). Cells were treated with 10 μM EdU for 1 hour before they were harvested, fixed, clicked, and analyzed for EdU incorporation. Unstained controls were included. Flow cytometry was performed using the CytoFLEX (Beckman Coulter). Cells positive for Alexa Fluor 647 were considered to be in S-phase and actively proliferating.

### Timelapse migration

Glass-bottom 96 well plates (Cellvis) were treated with L-poly-ornithine (Sigma) for 1 hour, followed by laminin coating. Cells were seeded on the plate at a density of 1.0-1.5×10^4^ cells/well and monitored for attachment before replacing media and placing in the LiveCyte (Phasefocus) cell imager. Cells were imaged every 20 minutes over a period of 24-48 hours. Quantification was performed by the LiveCyte software, with further graphical analysis being performed in GraphPad and R.

### Limiting dilution assay

hfNPCs or BTICs were plated in ultra low-attachment 96 well plates with 400, 200, 100, 50, 25, 12, 6, and 3 cells per well (12 wells per dilution). After 2 weeks, the fraction of wells containing neurospheres (tight, non-adherent masses > 50 μm in diameter) were recorded. The log of the fraction of non-responding wells was calculated and used for graphing results. Significance was determined through chi-squared test for stem cell frequency and pairwise comparisons for differences in stem cell frequencies using Extreme Limiting Dilution Assay (ELDA) webtool^115^.

### Recombinant CTSB experiments in GBM and NPC cell lines

CTSB (Sino Biological) was dissolved in base media and diluted to a final concentration of 100 nM, in the same range as previously published^116^. The dilution of CTSB was determined using a dilution curve in a cell viability assay compared to non-treated media control. For experiments, cells were treated for 48 hours in base media lacking growth factors unless otherwise noted.

### Senescence green and LysoTracker assay

hfNPCs were plated on a glass bottom 96 well plate and maintained in the presence or absence of CTSB for 48h. For an additional 1h, LysoTracker Red DND-99 (Invitrogen) was added to the cells at a final concentration of 50 nM. Cells were then washed twice with PBS and fixed with 2% PFA before being stained for β-galactosidase activity with the CellEvent™ Senescence Green Detection Kit (Invitrogen) per manufacturer instructions.

### Intraoperative LV-proximal and LV-distal sample collection

All de-identified samples and patient data were collected following Mayo Clinic Institutional Review Board (IRB) approved protocols and were collected only after obtaining informed consent from the patient. Brain tumor tissue was obtained during standard of care craniotomy for GBM resection by specialized neurosurgeons. Location of obtained biopsy samples was determined with intraoperative surgical navigation on the StealthStation S7 Surgical Navigation System (Medtronic PLC, Minneapolis, MN). Intraoperative screenshots of biopsy location were collected using the surgical navigation system and sample distance from the LV was determined by a physician using medical imaging software (Qreads Clinical Image Viewer, integrated to the Mayo Clinic EMR system). Tumors were considered LV-proximal when sample distance to LV was < 10 mm.

### RNA sequencing and analysis

Cells were grown in 25 cm^2^ flasks, transduced with lentivirus, and selected with puromycin until 80% confluent. Total RNA was isolated using TRIzol separation (ThemoFisher) combined with RNeasy column cleanup and DNase I treatment (Qiagen). Samples were diluted to a concentration of 60 ng/µL in nuclease-free water before shipping to the Mayo Clinic Genome Analysis Core. Samples were sequenced on the NovaSeq 6000 (Illumina) with paired end index read and the TruSeq v2 library.

The raw RNA sequencing paired-end reads for the samples were processed through the Mayo RNA-Seq bioinformatics pipeline, MAP-RSeq version 3.1.4^117^. Briefly, MAP-RSeq employs the very fast, accurate and splice-aware aligner, STAR^118^, to align reads to the reference human genome build hg38. The aligned reads are then processed through a variety of modules in a parallel fashion. Gene and exon expression quantification were performed using the Subread package^119^ to obtain both raw and normalized (FPKM) reads. Known and novel gene isoforms were assembled and quantified using StringTie^120^ to enable detection of alternative spliced isoforms. Finally, comprehensive analyses were run on the aligned reads to assess quality of the sequenced libraries.

Using the raw gene counts report from MAP-RSeq, genes differentially expressed between the groups were assessed using the bioinformatics package edgeR 2.6.2^121^. Genes found different between the groups are reported along with their magnitude of change (log2 scale) and their level of significance (q-value < 5%). Further analysis was performed using gene set enrichment analysis (GSEA) for hallmark signature gene sets to identify relevant biological pathways^122^.

### TCGA patient clustering

Of the 175 GBM RNA sequencing samples in TCGA_GBM database, 162 were from tumors and 161 had available survival data. The RNA expression counts (normalized FPKM values) for the 162 samples were downloaded. The RNA expression features were subset to 161 apoptosis related genes. Unsupervised clustering was performed on the 162 samples from the normalized FPKM values of the 161 apoptosis related genes using the R package Rphenograph (v0.99.1) algorithm with default options. Survival of the clusters was compared using the R package survminer (v0.4.9) and a p-value was computed using a log-rank test.

### Schematics and cartoons

All schematic and cartoon figures were created in Biorender.com.

### Statistical analysis

All data is represented as mean ± standard error of the mean (SEM) unless otherwise noted. Statistical analysis and graphical rendering were performed using GraphPad Prism® software and R Studio. Normal distribution of the data was evaluated using the Shapiro-Wilk normality test. For comparisons across two normally distributed groups, student’s t-test was performed. For matched patient sample immunohistochemistry, the paired sample t-test was performed. For comparisons among three normally distributed groups, one-way analysis of variance (ANOVA) with Tukey’s post-hoc correction was performed. For survival analysis, the Log-rank Mantel-Cox test was used and was corrected for multiple tests when more than two groups were analyzed. The level of significance was determined as p < 0.05. For all figures, * p < 0.05, ** p < 0.01, *** p < 0.001, **** p < 0.0001.

## Supplementary information

<u>Supplementary Figure 1 GBM BTICs have increased malignancy-associated phenotypes when co-cultured with hfNPCs</u>

(A) Viability assay over time of BTICs treated with conditioned medium from themselves or hfNPCs.

(B) Proliferation assay indicating percentage of Ki67+ BTICs when co-cultured with themselves or hfNPCs for 48h. Compared to NT conditions.

(C) Transwell migration assay of BTICs co-cultured with themselves or hfNPCs for 24h. Compared to NT conditions.

(D) Proliferation assay indicating percentage of Ki67+ hfNPCs when co-cultured with themselves or BTICs for 48h. Compared to NT conditions.

<u>Supplementary Figure 2 Nes-MetRS* mice effectively express GFP and MetRS* in NSCs when activated</u>

(A) IHC showing TAM-inducible GFP expression in Nestin+ SVZ cells. Scale bar = 50 µm.

(B) IHC showing GFP expression in rostral migratory stream (RMS). Scale bar = 50 µm.

(C) IHC showing GFP expression in subgranular zone (SGZ) of hippocampus. Scale bar = 25 µm.

(D) IHC showing TAMRA DBCO co-localization to GFP+ cells in the SVZ. Scale bar = 50 µm.

(E) Gel indicating increased TAMRA DBCO incorporation into SVZ lysate following TAM induction of NSCs.

(F) Silver stain of lysate immunoprecipitation showing pulldown enrichment in animals receiving both TAM and ANL.

<u>Supplementary Figure 3 Related to main figure 5</u>

(A) Percentage of Sox2+ cells in the SVZ in the presence of different shRNA GBMs proximal or distal to LV (n = 4-5).

(B) Percentage of Sox2+ cells in different shRNA GBMs proximal or distal to LV (n = 4-5).

(C) Percentage of TUNEL+ cells in different shRNA GBMs proximal or distal to LV (n = 4-5).

<u>Supplementary Figure 4 Soluble CTSB contributes to malignancy-associated phenotypes in BTICs</u>

(A) Cell viability measurements for NT and + recombinant CTSB at 48h.

(B) Proliferation assay indicating percentage of EdU+ cells in NT and + recombinant CTSB.

(C) Transwell migration of NT and + recombinant CTSB.

(D) Total distance traveled by NT and + recombinant CTSB in timelapse migration.

(E) Distance from origin traveled by NT and + recombinant CTSB in timelapse migration.

(F) Representative plots of migration from origin for NT and + recombinant CTSB in BTIC line QNS120.

<u>Supplementary Figure 5 Apoptosis gene signature expression by TCGA GBM patient clusters</u>

Heatmap showing relative gene expression for genes in the apoptosis hallmark gene expression set from MSigDB in TCGA GBM patients by cluster.

<u>Supplementary Table: Patient information for intraoperative specimen collection</u>

