## Supplementary figures and images for "Cell-specific crosstalk proteomics reveals cathepsin B signaling as a driver of glioblastoma malignancy near the subventricular zone"

### Supplemental figure 1

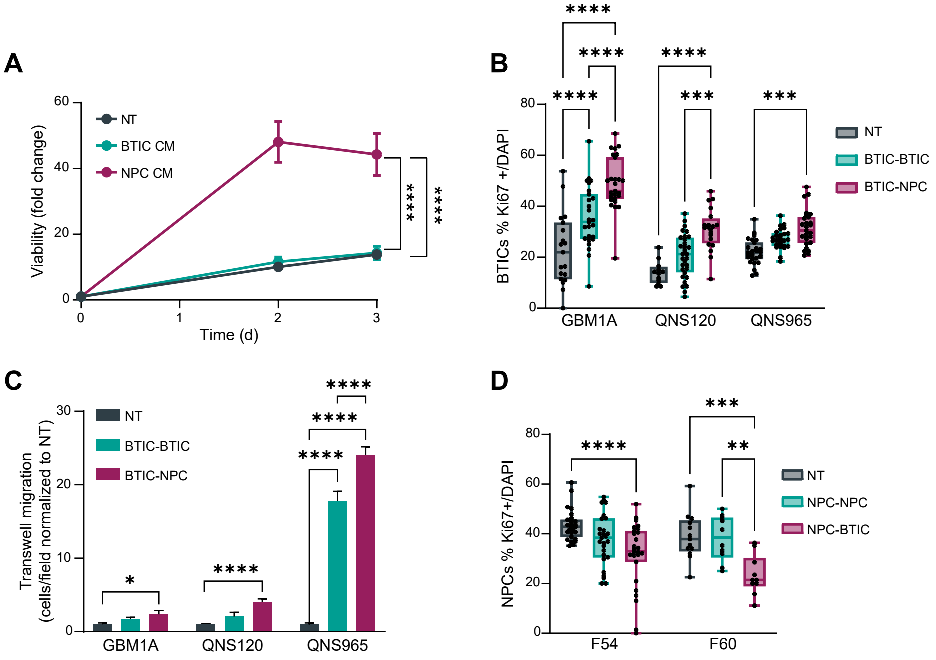

### Supplemental figure 2

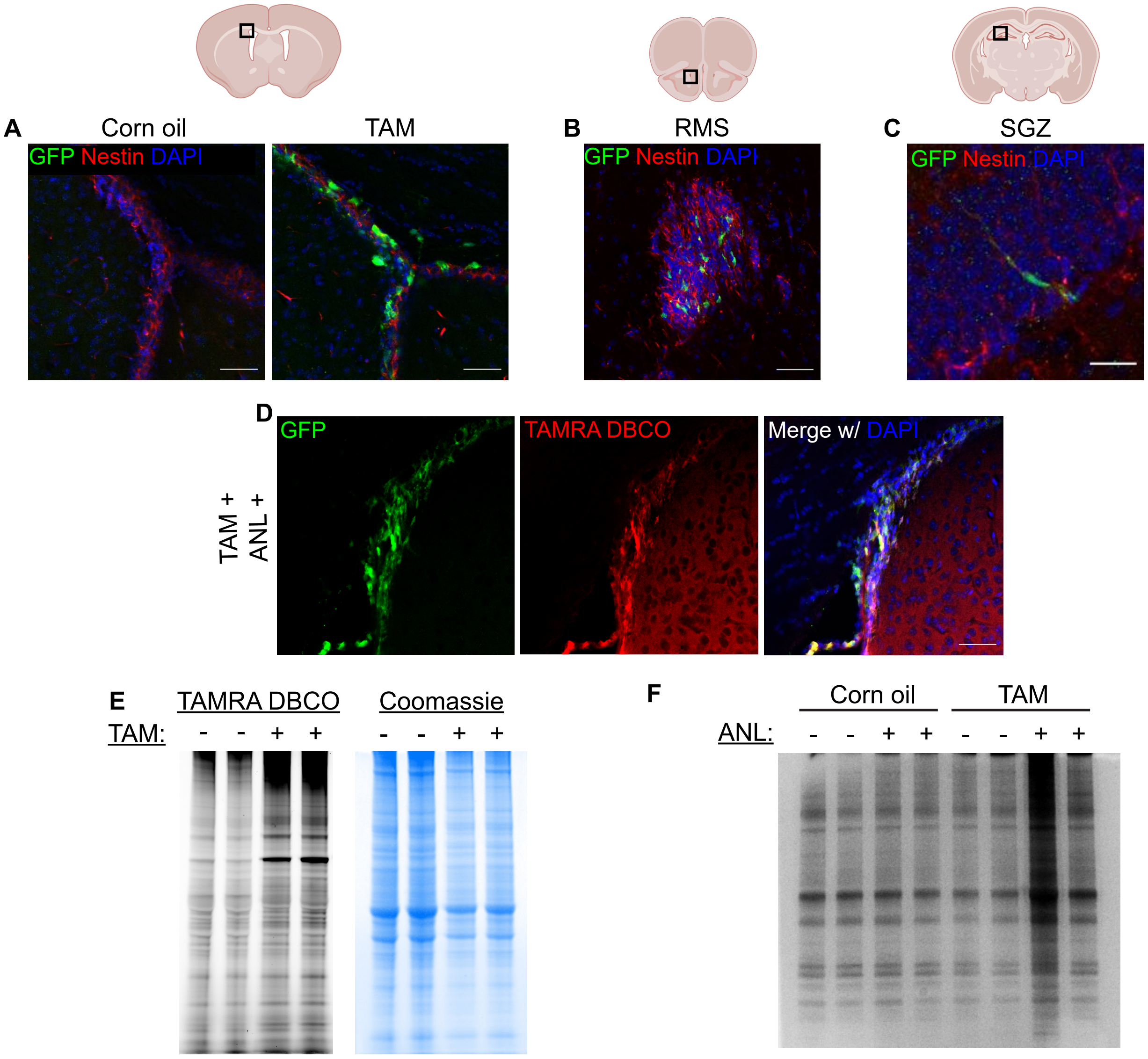

### Supplemental figure 3

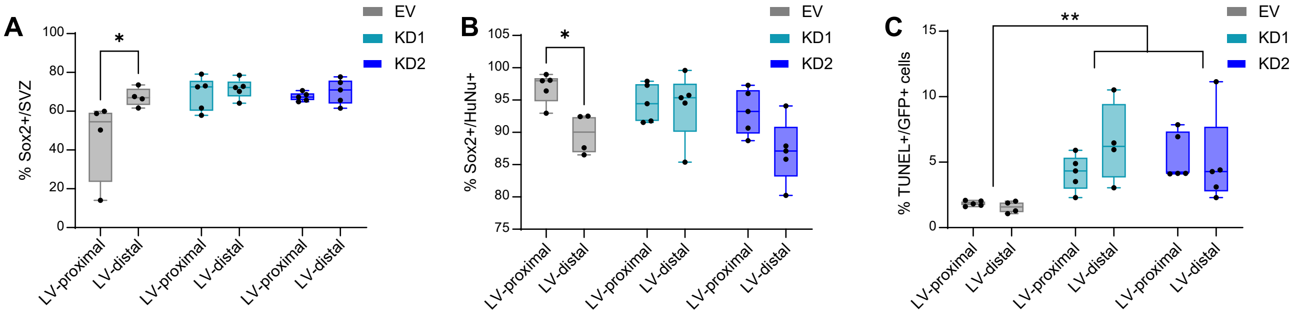

### Supplemental figure 4

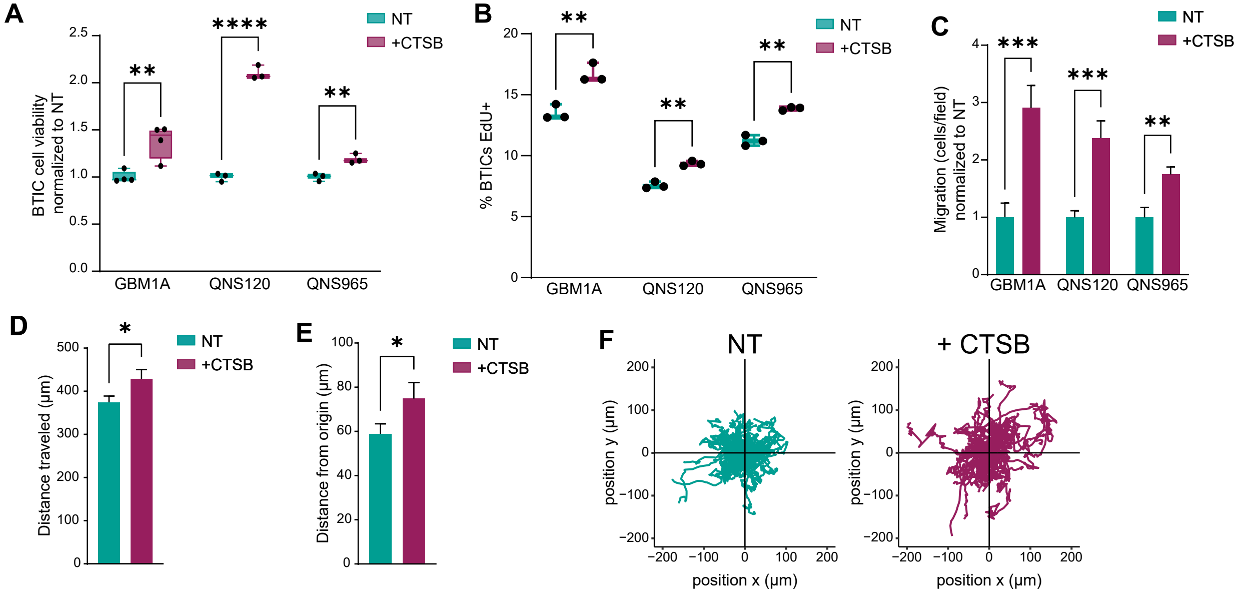

### Supplemental figure 5

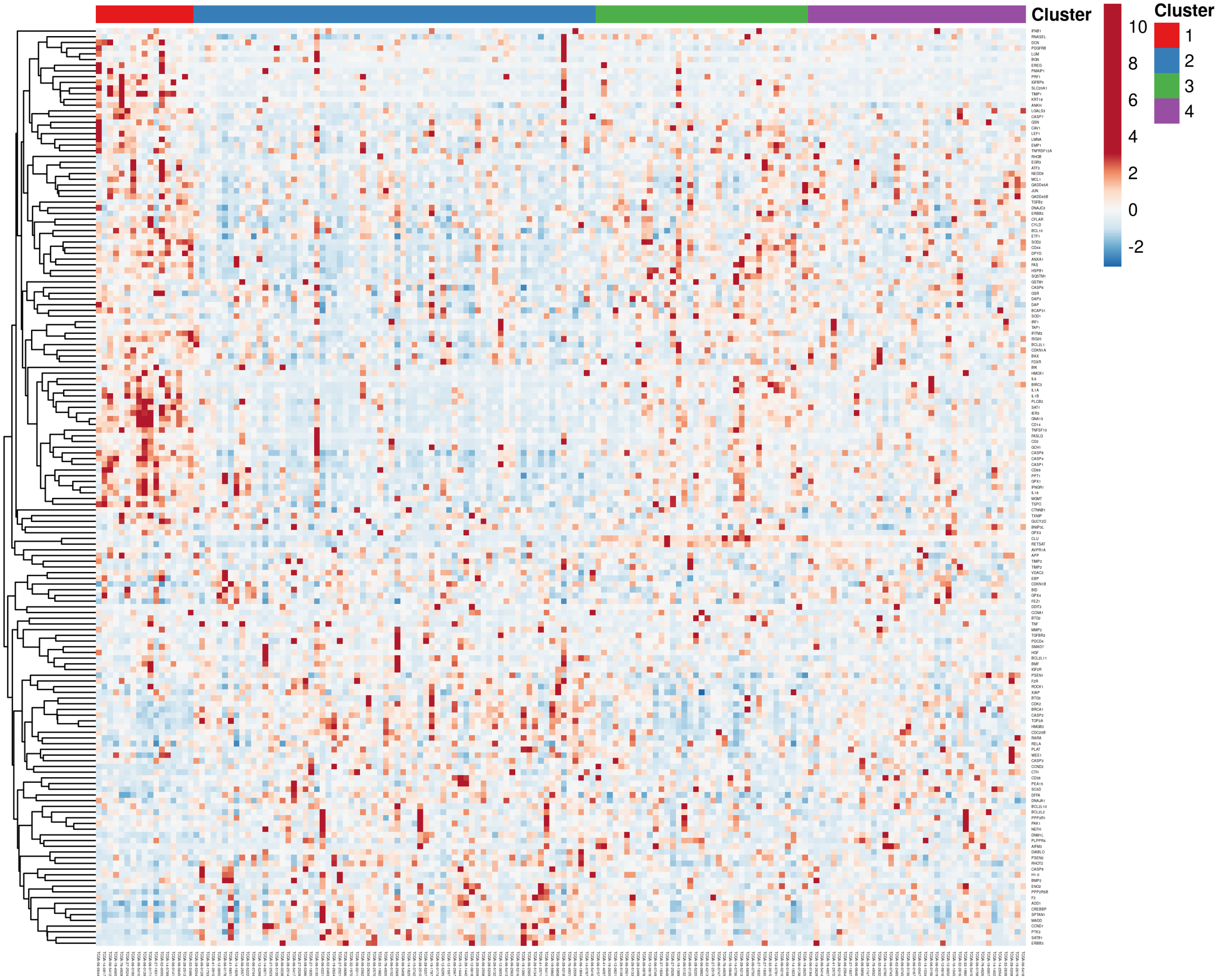
